# The myocardium utilizes Pdgfra-PI3K signaling to steer towards the midline during heart tube formation

**DOI:** 10.1101/2023.01.03.522612

**Authors:** Rabina Shrestha, Tess McCann, Harini Saravanan, Jaret Lieberth, Prashanna Koirala, Joshua Bloomekatz

## Abstract

Coordinated cell movement is a fundamental process in organ formation. During heart development, bilateral myocardial precursors collectively move towards the midline (cardiac fusion) to form the primitive heart tube. Along with extrinsic influences such as the adjacent anterior endoderm which are known to be required for cardiac fusion, we previously showed that the platelet-derived growth factor receptor alpha (Pdgfra) is also required. However, an intrinsic mechanism that regulates myocardial movement remains to be elucidated. Here, we uncover an essential intrinsic role in the myocardium for the phosphoinositide 3-kinase (PI3K) intracellular signaling pathway in directing myocardial movement towards the midline. *In vivo* imaging reveals that in PI3K-inhibited zebrafish embryos myocardial movements are misdirected and slower, while midline-oriented dynamic myocardial membrane protrusions become unpolarized. Moreover, PI3K activity is dependent on and genetically interacts with Pdgfra to regulate myocardial movement. Together our findings reveal an intrinsic myocardial steering mechanism that responds to extrinsic cues during the initiation of cardiac development.

## Introduction

During organogenesis, cell progenitor populations often need to move from their origin of specification to a new location in order to form a functional organ. Deficient or inappropriate movement can underlie congenital defects and disease. Directing these movements can involve extrinsic factors such as chemical and mechanical cues from neighboring tissues and the local environment as well as intrinsic mechanisms such as intracellular signaling and polarized protrusions (1). Progenitor cell movement occurs during cardiac development, where myocardial cells are specified bilaterally on either side of the embryo (2). To form a single heart that is centrally located, these bilateral populations must move to the midline and merge (3, 4). As they move, myocardial cells undergo a mesenchymal-to-epithelial (MET) transition forming intercellular junctions and subsequently moving together as an epithelial collective (5-9). This process is known as cardiac fusion and occurs in all vertebrates (10, 11).

External influence from the adjacent endoderm is essential for the collective movement of myocardial cells towards the midline. Mutations in zebrafish and mice which inhibit endoderm specification or disrupt endoderm morphogenesis result in cardia bifida – a phenotype in which the bilateral myocardial populations fail to merge (9, 12-21). Similar phenotypes also occur in chicks and rats when the endoderm is mechanically disrupted (22-24). Studies simultaneously observing endoderm and myocardial movement have found a correlation between the movements of these two tissues, suggesting a model in which the endoderm provides the mechanical force that pulls myocardial cells towards the midline (24-27). Yet, these correlations do not occur at all stages of cardiac fusion, indicating that myocardial cells may also use intrinsic mechanisms to actively move towards the midline. Indeed, recent studies revealing a role for the receptor tyrosine kinase, platelet-derived growth factor receptor alpha (Pdgfra) in the movement of myocardial cells have suggested a paracrine chemotaxis model, in which the myocardium senses chemokine signals from the endoderm and responds to them (28). However, the existence and identity of these intrinsic myocardial mechanisms remain to be fully elucidated.

We have sought to identify the intracellular pathways downstream of Pdgfra that regulate the collective movement of the myocardium. The phosphoinositide 3-kinase (PI3K) pathway is known as an intracellular signaling mediator of receptor tyrosine kinases (e.g. Pdgfra). PI3K phosphorylates phosphatidylinositol (4,5)-bisphosphate (PIP2) to create phosphatidylinositol (3,4,5)-trisphosphate (PIP3), a regulator of cellular processes such as proliferation and cell migration (29). The PI3K pathway has been shown to be important for both individualistic cell migration such as in Dictyostelium and neutrophils (30, 31) as well as collective cell migration such as in the movement of border cells in Drosophila and the movement of the anterior visceral endoderm during mouse gastrulation (32, 33).

Using the advantages of external development and ease of live-imaging in the zebrafish model system (34), our studies reveal that myocardial PI3K signaling is required for proper directional movement towards the midline during cardiac fusion. In particular we find that inhibition of the PI3K pathway, throughout the embryo or only in the myocardium, results in bilateral cardiomyocyte populations that fail to reach the midline (cardia bifida) or have only partially merged by the time wild-type myocardial cells are fully merged. High-resolution live imaging in combination with mosaic labeling further reveals that the orientation of myocardial membrane protrusions during cardiac fusion is dependent on PI3K signaling. Furthermore, we find that PI3K signaling and Pdgfra genetically interact to facilitate cardiac fusion. Altogether our work supports a model by which intrinsic Pdgfra-PI3K signaling regulates the formation of membrane protrusions that facilitate the collective movement of the myocardium towards the midline. Insight into the balance of extrinsic and intrinsic influences for directing collective movement of myocardial cells has implications for understanding a wide set of congenital and environmental cardiac defects as well as the pathogenic mechanisms of diseases broadly associated with collective movement.

## Results

### The PI3K pathway is required for proper cardiac fusion

In a search for intracellular signaling pathways that are important for cardiac fusion we examined the role of the phosphoinositide 3-kinase (PI3K) signaling pathway, by pharmacological inhibition of PI3K with LY294002 (LY) (35). Treatments were started at the bud-stage (10 hours post-fertilization -hpf), in order to exclude effects on mesodermal cells during gastrulation (36). In wild-type or DMSO-treated embryos, bilateral myocardial populations move towards the midline and merge to form a ring structure between 20-21 hpf, which corresponds to the 20-22 somite stage (s) (Fig. 1A, A’, F). However, in LY-treated embryos myocardial movement is disrupted and the bilateral myocardial populations fail to properly merge by 22s (Fig. 1B, B’, F, Suppl. Fig. 1A-C, M). To rule out possible off-target phenotypic artifacts of LY (37), we exposed bud stage embryos to two other PI3K inhibitors, Dactolisib (Dac) or Pictilisib (Pic) (38, 39). Exposure to either of these inhibitors also causes cardiac fusion defects (Fig. 1C, C’, F, G, Suppl. Fig. 1D-F, N; Fig. 1D, D’, F, G, Suppl. Fig. 1G-I, O, respectively), as does the mRNA injection of a truncated form of p85 (Fig. 1E, E’, F, G, Suppl. Fig. 1J-L, P), which acts as a dominant negative inhibitor of PI3K (dnPI3K) activity (40). Furthermore, to ensure our analysis was not complicated by a developmental delay, we used developmentally stage-matched embryos (somite stage) rather than time-matched embryos (hours post-fertilization; hpf) to assess cardiac fusion phenotypes (see Suppl. Fig. 2 for embryos analyzed at 20 hpf).

**Fig. 1.**
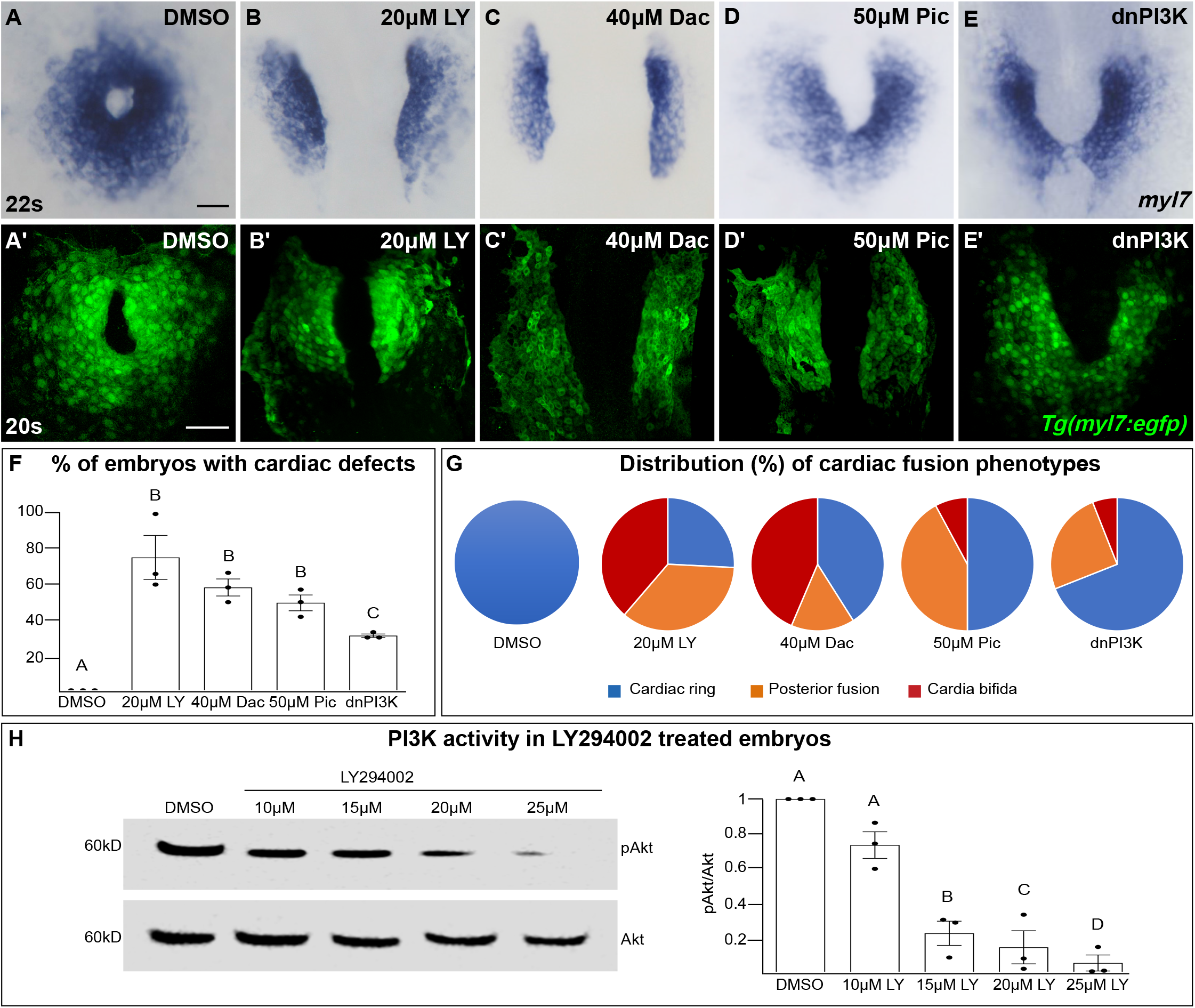
The PI3K pathway is required for cardiac fusion. **A-E** Dorsal views, anterior to the top, of the myocardium labeled with *myl7* (A-E) at 22 somite stage (s) or *Tg(myl7:egfp)* (A’-E’) at 20s. In contrast to a ring of myocardial cells in DMSO-treated embryos (A, A’), in embryos treated with PI3K inhibitors LY294002 (LY, B, B’), Dactolisib (Dac, C, C’), or Pictilisib (Pic, D, D’) at bud stage or injected with *dnPI3K* mRNA (750pg) at the one-cell stage (E, E’) cardiac fusion fails to occur properly with embryos displaying either cardia bifida (B, C) or fusion only at the posterior end (D, E). **F, G** Graphs depict the percentage (F) and range (G) of cardiac fusion defects in control and PI3K-inhibited embryos. Dots represent the percent of embryos with cardiac defects per biological replicate. Total embryos analyzed n = 37 (DMSO), 31 (20µM LY), 39 (40µM Dac), 38 (50µM Pic), 86 (*dnPI3K*). Blue – cardiac ring/normal; Orange – fusion only at posterior end/mild phenotype, Red – cardia bifida/severe phenotype. **H** Representative immunoblot and ratiometric analysis of phosphorylated Akt (pAkt) to Akt protein levels in DMSO and LY treated embryos reveals a dose-dependent decrease in PI3K activation. Bar graphs indicate mean ± SEM, dots indicate pAKT/AKT ratio per biological replicate, normalized to DMSO. Three biological replicates per treatment. One-Way ANOVA tests – letter changes indicate differences of p < 0.05 (F, H). Scale bars, 40 µm (A-E), 42 µm (A’-E’). Raw data and full p-values included in the source file.

We also examined the morphology of the cardiac ring in PI3K-inhibited embryos and cellular processes known to be regulated by PI3K signaling. During the later stages of cardiac fusion as part of the subduction process, medial myocardial cells form a contiguous second dorsal layer (26) and develop epithelial polarity in which intercellular junction proteins such as ZO1 are localized to the outer-edge of the myocardium (5, 6) (Suppl. Fig. 3A-C). In PI3K-inhibited embryos, we found that myocardial cells form this second dorsal layer however, the localization of polarity markers and the tissue organization can appear mildly disorganized (Suppl. Fig. 3D-F). Furthermore, the PI3K signaling pathway is known to promote cell proliferation and cell survival (29) however, we did not find a difference in the number of cardiomyocytes in DMSO-or LY-treated embryos at 20s (Suppl. Fig. 3G-I). Similarly, no apoptotic cardiomyocytes were observed in DMSO-nor in 20 µM LY-treated embryos (n = 17, 19 embryos, respectively from 3 biological replicates). Apoptotic cardiomyocytes were however observed in DNAse-treated controls. These experiments reveal that PI3K signaling is required for proper cardiac fusion.

### The extent and duration of PI3K inhibition determines the penetrance and severity of cardiac fusion defects

PI3K-inhibited embryos display cardiac phenotypes at 22s that range from severe, in which the myocardial populations remain entirely separate (cardia bifida) (Fig. 1G – red; examples -Suppl. Fig 1C, F, I, L), to more mildly affected hearts in which the myocardial populations form a U-shaped structure, having merged at the posterior but not anterior end (Fig. 1G – orange; examples Suppl. Fig. 1B, E, H, K). A subset of the PI3K-inhibited embryos also appear phenotypically normal (∼25% for 20 µM LY, Fig. 1F, G) indicating incomplete penetrance. Increasing concentrations of PI3K inhibitor or dnPI3K mRNA increases the severity and penetrance of these phenotypes in a dose-dependent manner (Suppl. Fig. 1.). Similarly, we confirmed that LY inhibits PI3K activity in a dose-dependent manner, as measured by the ratio of phosphorylated AKT (pAKT) to AKT (Fig. 1H). AKT is phosphorylated as a direct consequence of PI3K activity (41). Thus, the severity and penetrance of cardiac fusion defects depends on the efficacy of PI3K inhibition.

Since differing modes of movement (9) as well as cellular processes such as MET (5, 6) and subduction (26) occur at different times during cardiac fusion, we also evaluated the developmental stages over which PI3K signaling is required. Short exposures (<3 hours) just prior to 22s or starting at bud stage had no effect on cardiac fusion. However, progressively longer times of exposure ending at 22s or starting at bud stage result in correspondingly more severe phenotypes and higher penetrance (Fig. 2A, B). These addition and wash-out experiments indicate that both the severity and penetrance of cardiac fusion phenotypes correlate with the duration of LY-incubation and not a specific developmental stage inside the 3-20s window. Thus, myocardial movement is responsive to both the levels and duration of PI3K signaling throughout cardiac fusion.

**Fig 2:**
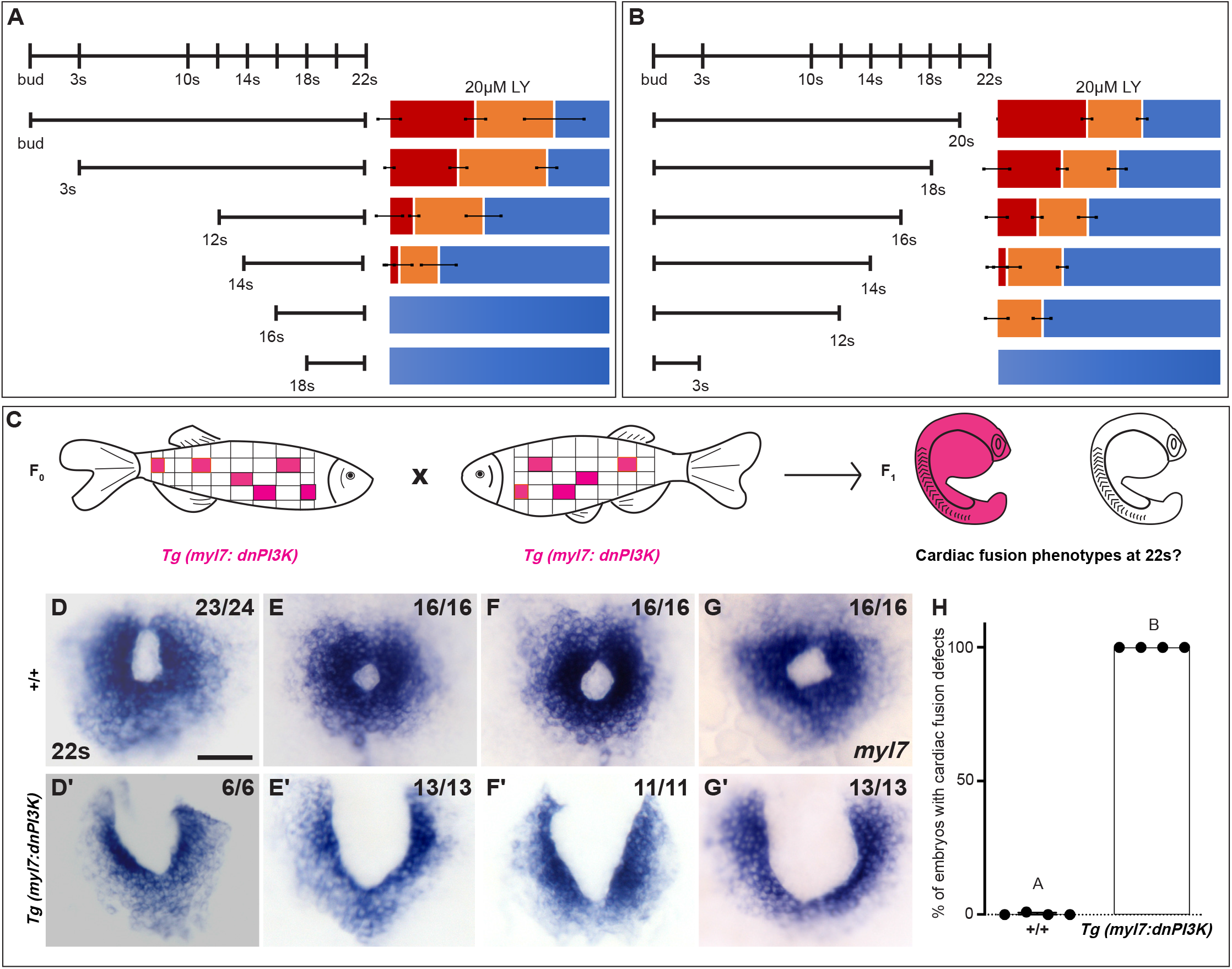
PI3K is required in the myocardium throughout cardiac fusion. **A, B** Graphical representation of the PI3K inhibitor addition (A) and wash-out (B) experiments used to determine the developmental stage over which PI3K is required. In (A) LY is added to embryos at different developmental stages and incubated until 22s, when cardiac fusion is assessed. In (B) LY is added at bud stage and washed-out at different developmental stages, after which embryos are incubated in normal media till 22s, when cardiac fusion is assessed. Bar graphs indicate the average proportion of embryos displaying different phenotypes. Blue – cardiac ring/normal; Orange – fusion only at posterior end/mild phenotype, Red – cardia bifida/severe phenotype. n = 45 embryos per treatment condition from three biological replicates. **C** Schematic outlines experimental design to test requirement for PI3K in the myocardium. Pink – cells with the *Tg(myl7:dnPI3K)* transgene. F0 animals are mosaic for the transgene, while all cells in F1 embryos either have the transgene (pink) or do not (clear). The *myl7* promoter restricts *dnPI3K* expression to the myocardium in *Tg(myl7:dnPI3K)* embryos. **D-G** Dorsal view of the myocardium labeled with *myl7* in embryos at 22s from 4 different founder pairs (D-D’, E-E’, F-F’, G-G’). F1 embryos without the *Tg(myl7:dnPI3K)* transgene (as determined by genotyping) display normal cardiac fusion (D-G, n = 23/24, 16/16, 16/16, 16/16, per founder pair), while F1 siblings with the *Tg(myl7:dnPI3K)* transgene display cardiac fusion defects (D’-G’, n = 6/6, 13/13, 11/11, 13/13), indicating that PI3K signaling is required in myocardial cells. **H** Graph indicating the average % of wild-type and *Tg(myl7:dnPI3K)+* embryos with cardiac fusion defects. Letter difference indicates a significant Fisher’s exact test p = 5.56 × 10^−31^. Scale bar, 40µm.

### PI3K signaling is required in the myocardium for proper cardiac fusion

Mutations affecting the specification or morphology of the anterior endoderm result in myocardial movement defects (13, 19, 42, 43), revealing a non-autonomous role for the anterior endoderm in cardiac fusion. However, when PI3K signaling is inhibited with 15 or 25 µM LY starting at bud stage we did not observe differences in the expression of endoderm markers such as *axial/foxa2* or *Tg(sox17:egfp)* compared to DMSO-treated embryos (Suppl. Fig 3J-L, N-P). Additionally, the overall morphology of the anterior endoderm appeared intact and the average anterior endoderm width was similar between PI3K-inhibited and DMSO-treated embryos (Suppl. Fig 3M, Q).

To determine if PI3K signaling is specifically required within the myocardium, as opposed to the endoderm, we created a myocardial-specific dominant negative transgenic construct, *Tg(myl7:dnPI3K)*. Our experimental design is outlined in Fig. 2C. In F1 embryos at 22s we observed embryos with normal cardiac rings and embryos with cardiac fusion defects (Fig. 2D-G’). Genotyping revealed that F1 embryos with normal cardiac rings (Fig. 2D-G) did not have the transgene (n = 71/71), while almost all sibling embryos with cardiac fusion defects (Fig. 2D’-G’) were positive for the *Tg(myl7:dnPI3K)* transgene (n=40/41). And all embryos with the *Tg(myl7:dnPI3K)* transgene have a cardiac fusion defect (Fig. 2H). (F1 embryos from 4 independent founder pairs were analyzed since stable transgenics could not be propagated due to loss of viability, likely due to a requirement for PI3K signaling in cardiac contraction at later stages (44)). Statistical analysis reveals that the *Tg(myl7:dnPI3K)* transgene is significantly associated with a cardiac fusion defect (Fisher’s test p = 5.56 × 10^−31^), indicating that PI3K signaling acts in the myocardium to regulate its movement during cardiac fusion.

### PI3K signaling is responsible for the steering and velocity of myocardial movements during cardiac fusion

Our analysis points to a role for PI3K signaling in the movement of myocardial cells. To identify the properties of myocardial movement regulated by PI3K signaling, we analyzed myocardial movement by performing *in vivo* time-lapse imaging with the *Tg(myl7:egfp)* transgene, which labels myocardial cells. A time-series using *hand2* expression to compare myocardial movement in DMSO-treated and PI3K-inhibited embryos reveals dramatic differences in myocardial movement beginning after 12s (Suppl. Fig. 4). We thus focused our time-lapse imaging on the 14-20s developmental window. In time-lapse movies of DMSO-treated embryos, myocardial cells display coherent medially directed movement (Fig. 3A-B, E, Suppl. Fig. 5A-A’’’, Video-1) with an average velocity of 0.2334 ± 0.007 microns/min, which is consistent with previous studies (9, 28). In PI3K-inhibited embryos myocardial cells also display coherent, coordinated movement and do move in the general direction of the midline, however they make dramatically less progress (Fig. 3C-D, Suppl. Fig. 5B-B’’’, Video-2). Quantitative analysis of these myocardial cell tracks reveals that myocardial cells are slower (0.1879 ± 0.008 microns/min) and less efficient. Therefore, myocardial cells ultimately move less in LY-inhibited embryos compared to DMSO-treated embryos (Fig. 3E, F, Video-3). Differences in velocity occur throughout cardiac fusion (Suppl. Fig. 5C). However, the most dramatic difference between PI3K-inhibited and DMSO-treated myocardial cells is in the direction of their movement. Tracks of myocardial cells in DMSO-treated embryos are predominately oriented in a medial direction (average of 31.1 ± 1.65 degrees), while tracks in LY-treated embryos are mostly oriented in an angular anterior direction (60.6 degrees ± 1.73, p-value = 2.77 × 10^−12^, Fig. 3G, H). Differences in directional movement occur mainly in the early stages of cardiac fusion when wild-type myocardial movement is mostly medial (Suppl. Fig. 5D). Together this analysis of myocardial cell tracks suggests that PI3K signaling is responsible for both steering and propelling myocardial cells towards the midline.

**Fig 3:**
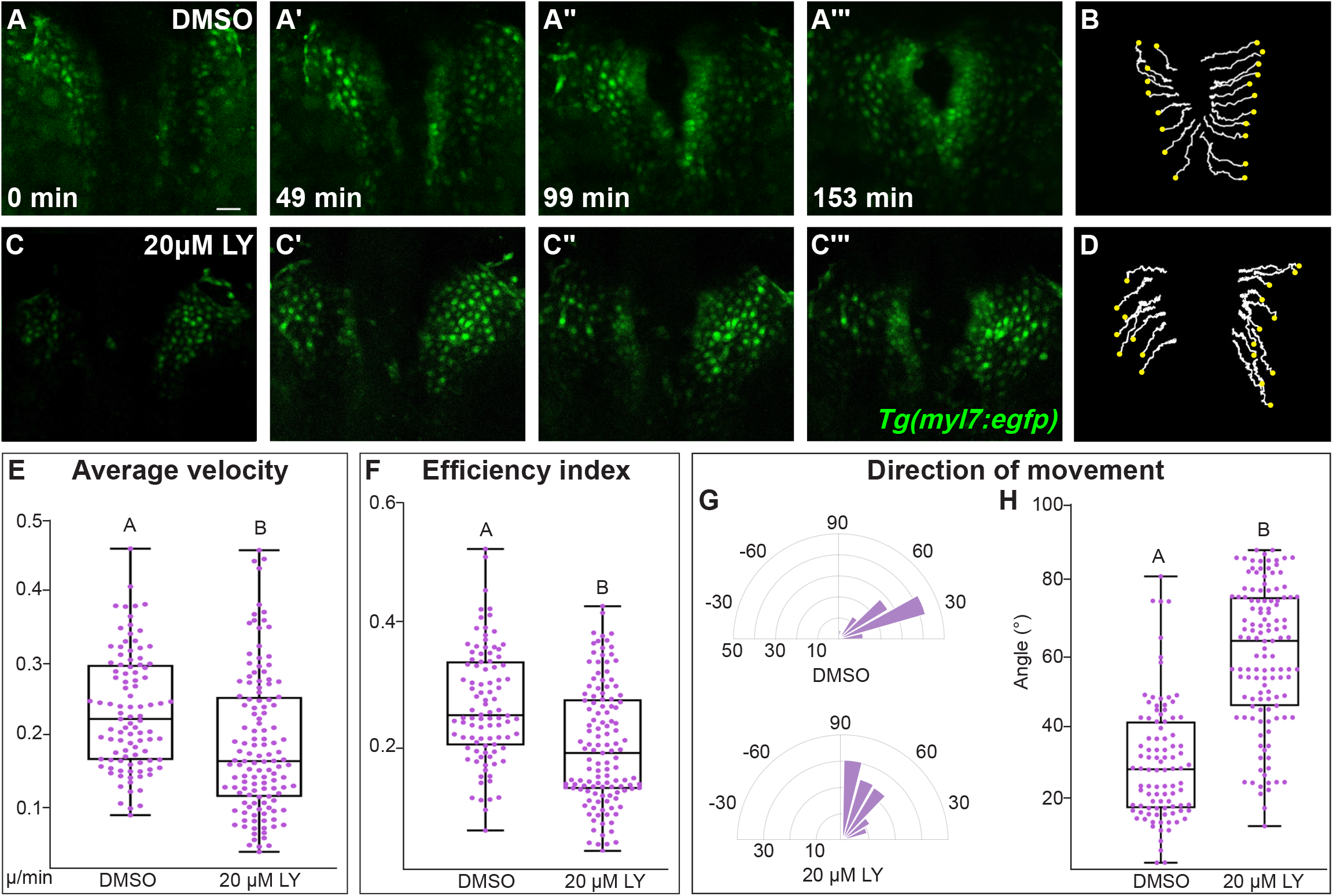
PI3K signaling regulates the medial movement and velocity of the myocardium during cardiac fusion. **A-D** Time points from a representative time-lapse video of myocardial cells visualized with the *Tg(myl7:egfp)* transgene in embryos treated with DMSO (A, B, Video-1) or 20µM LY (C, D, Video-2) from bud -22s. 3D reconstructions of confocal slices (A, C) reveal the changes in conformation and location of the myocardium at 3 major stages of cardiac fusion: early medial movement towards the embryonic midline (A-A’, C-C’), posterior merging of bilateral populations (A’’, C’’) and anterior merging to form a ring (A’’’, C’’’). Representative tracks (B, D) show the paths of a subset of myocardial cells over ∼2.5 hr timelapse. Yellow dots indicate the starting point of each track. **E-H** Graphs depict box-whisker plots of the velocity (E), efficiency index (F) and angle of movement (H) of myocardial cells. The direction of movement is visualized by rose plots (G). Myocardial cells in PI3K-inhibited (LY-treated) embryos show an overall direction of movement that is angular (60-90 degrees) and is slower than in DMSO-treated embryos. Scale bar, 60µm. 96 and 125 cells were analyzed from five DMSO- and six 20μM LY-treated embryos, respectively. Two sample t-test, letter change indicates p < 0.05. Raw data and full p-values included in the source file.

### Myocardial membrane protrusions are medially polarized by PI3K signaling

The role of PI3K signaling in regulating the polarity of migratory protrusions in the dorsal epithelium in Drosophila and prechordal plate in zebrafish (36, 45) along with previous reports of the existence of myocardial membrane protrusions (7, 26), led us to next look for these protrusions in myocardial cells during cardiac fusion and to examine if they are disrupted in PI3K-inhibited embryos. To visualize membrane protrusions in the myocardium, we performed *in vivo* time-lapse imaging during cardiac fusion of embryos injected with *myl7:lck-egfp* plasmid DNA in order to mosaically label the plasma membrane of myocardial cells. Despite myocardial cells being connected via intercellular junctions (5, 28), we observed that the lateral edges of myocardial cells in wild-type/DMSO-treated embryos are highly dynamic; transitioning from appearing smooth and coherent to undulating and extending finger-like membrane protrusions away from the cell (Fig. 4A-A’’’’, Video-4). These protrusions are dynamic, actively extending and retracting, and are prevalent occurring on average 20.3 ± 6.7 times per hour per cell and lasting for an average of 2.3 ± 0.6 mins (Fig. 4A). In LY-treated embryos we observed similar membrane protrusions extending from myocardial cells (Fig. 4B,Video-4), which occur at a similar rate (17 ± 7.4 per hour per cell, p-value = 0.36), but with slightly longer persistence (3.23 ± 0.84 mins, p-value = 0.008).

We further observed that in DMSO-treated embryos membrane protrusions occur predominantly in the medial direction (77.25 ± 21.76% of protrusions were in the forward direction, Fig. 4A-A’’’’, C, D), suggesting an association with the medial movement of the myocardial tissue. In contrast, in LY-treated embryos myocardial membrane protrusions do not display the same medial polarity, instead extending from all sides of a myocardial cell equally (only 46 ± 11.6% of protrusion were in the forward direction, Fig 4B-B’’’’, C, D). The finding that myocardial membrane protrusions are medially polarized in wild-type embryos but not in PI3K-inhibited embryos where myocardial cells are misdirected and slower to reach the midline suggests that PI3K signaling helps to steer and propel myocardial cells towards the midline through the polarization of these active protrusions.

**Fig 4:**
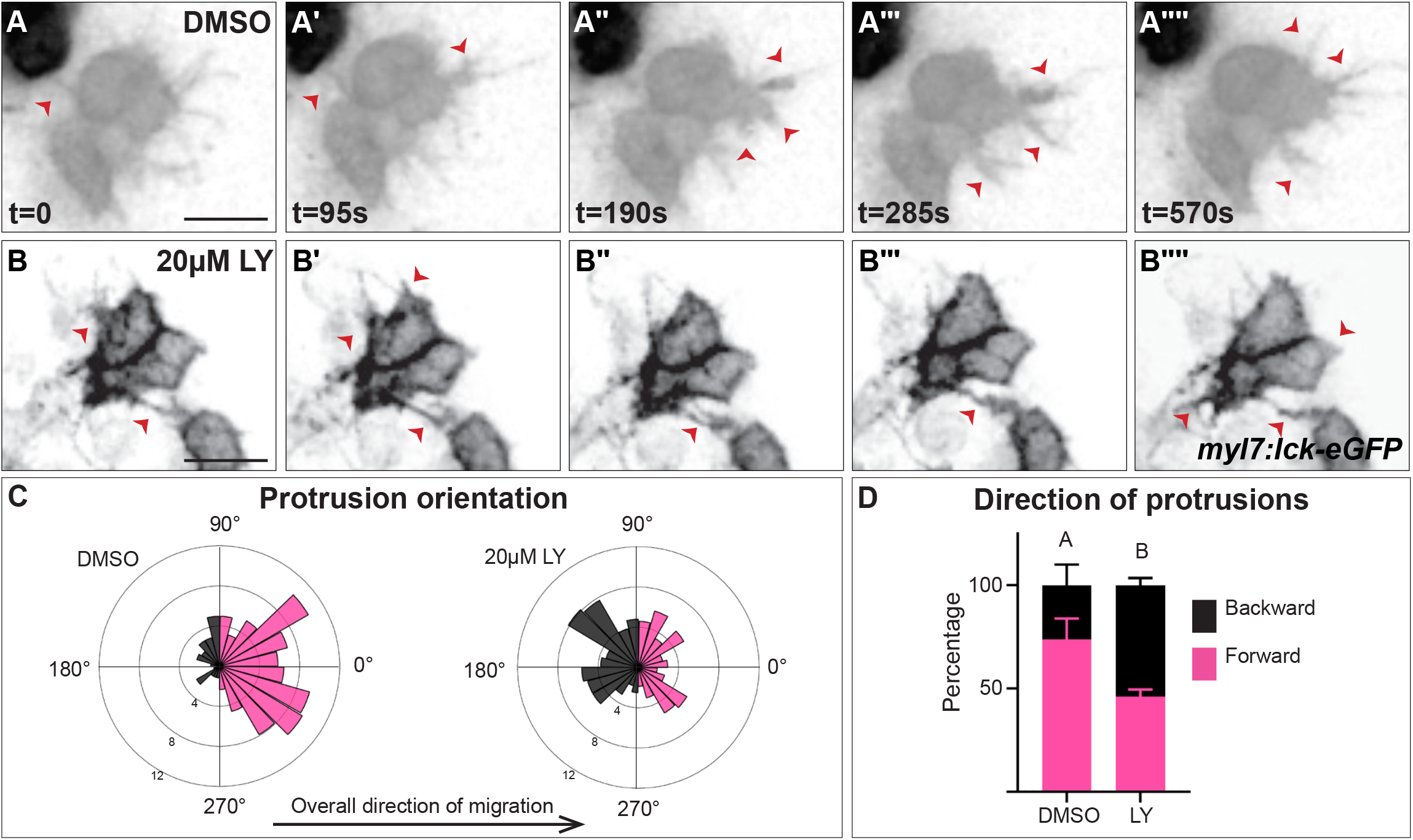
Myocardial membrane protrusions are misdirected in PI3K-inhibited embryos. **A-B’’’’** timepoints from representative timelapse videos (see Video-4) of myocardial cells whose membrane has been labeled with *myl7*:lck-eGFP (black), medial to the right, in a DMSO-(A-A’’’’) or a 20µM LY-(B-B’’’’) treated embryo. Red arrowheads indicate representative protrusions, which are oriented medially, coincident with the direction of movement in DMSO-treated embryos (A-A’’’’) but are oriented in all directions in LY-treated embryos (B-B’’’’). **C, D** Rose (C) and Bar (D) graphs displaying the orientation of membrane protrusions in DMSO-(left) or LY-(right) treated embryos. The length of each radial bar in (C) represents the percentage of protrusions in each bin. Bar graph displays the total percentage of forward or backward protrusions. Forward protrusions: 270-90 degrees, pink. Backward protrusions: 90-270 degrees, black. n = 425 protrusions from 11 cells in 5 embryos (DMSO), and 480 protrusions from 11 cells in 4 embryos (20µM LY). Fisher’s exact test, P value < 0.0001. Error bars, mean ± SEM. Scale bar, 30 µm. Raw data and full p-values included in the source file.

### PI3K signaling is regulated by Pdgfra during cardiac fusion

The improperly directed myocardial cells in PI3K-inhibited embryos (Fig. 3) are reminiscent of the steering defects observed in *pdgfra* mutant embryos (28). This similarity led us to investigate whether Pdgfra activates PI3K signaling to regulate myocardial movement. We found that PI3K activity as measured by the ratio of phospho-AKT to AKT levels (41), is severely diminished in *pdgfra* mutant embryos during cardiac fusion (Fig. 5A). Conversely, when Pdgfra activity is increased during cardiac fusion through the over-expression of *pdgf-aa*, PI3K activity is up-regulated (Fig. 5B).

**Fig 5:**
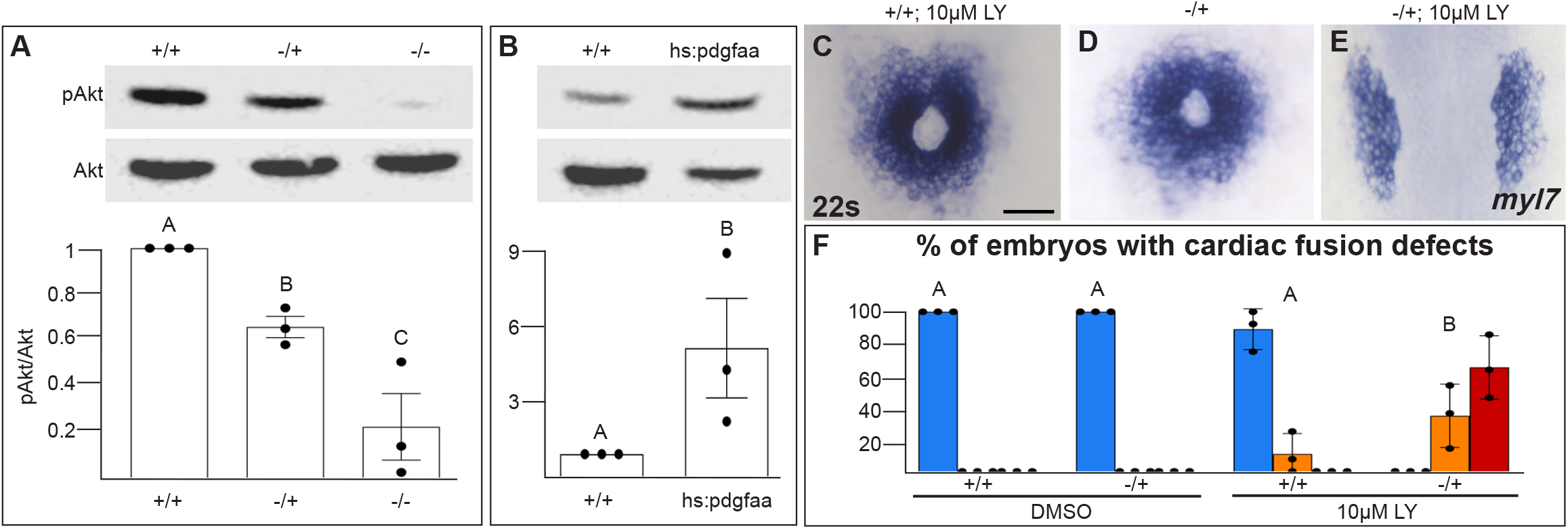
Pdgfra activates and genetically interacts with PI3K signaling to regulate cardiac fusion. **A, B** Immunoblot and ratiometric analysis of phosphorylated Akt (pAkt) compared to total Akt levels reveals reduced pAkt levels in loss-of-function *pdgfra*^*sk16*^ heterozygous (-/+) or homozygous (-/-) mutant embryos at 22s (A), and elevated pAkt levels at 22s when PDGF signaling is activated with the *hs:pdgfaa* transgene (B). Bar graphs display averages from three separate experiments. **C-E** Dorsal views, anterior to the top, of the myocardium labeled with *myl7* at 22s. In contrast to a normal ring of myocardial cells in wild-type embryos treated with 10µM LY starting at bud stage (C) or *pdgfra* heterozygous embryos (D), when *pdgfra* heterozygous mutants are exposed to 10µM LY, cardiac fusion is defective with embryos displaying severe phenotypes such as cardia bifida (E). **F** Bar graph depicts the average distribution of cardiac fusion defects in DMSO-treated wild-type and *pdgfra* heterozygous mutants as well as 10µM LY-treated wild-type and *pdgfra* heterozygous mutant embryos. The total number of embryos examined over three separate biological replicates are 47 (DMSO, +/+), 25 (DMSO, -/+), 36 (10µM LY, +/+), and 31 (10µM LY, -/+). Blue - cardiac ring/normal; Orange - fusion only at posterior end/mild, Red - cardia bifida/severe. Bar graphs, mean ± SEM. One-way ANOVA (A, C) or Student’s T-test (B), letter change indicates p < 0.05. Scale bar, 60µm. Raw data and full p-values included in the source file.

To determine if Pdgfra’s influence on PI3K activity is important for myocardial movement towards the midline, we examined whether these two genes genetically interact while regulating cardiac fusion. When *pdgfra* heterozygous mutant embryos are exposed to DMSO cardiac fusion occurs normally (Fig. 5D, F), even though there is a small reduction in PI3K activity (Fig. 5A). When wild-type embryos are exposed to 10µM LY, PI3K activity is modestly reduced (Fig. 1H) and a small percent of embryos display mild cardiac fusion defects (Average of 10.9 ± 7.39% of 10µM LY-treated embryos display mild U-shaped cardiac fusion defects, n= 36, 3 replicates, Fig 5C, F). However, when *pdgfra* heterozygous mutant embryos are exposed to 10µM LY, there is a synergistic increase in both the severity and penetrance of cardiac fusion defects. 100% of *pdgfra* heterozygous embryos exposed to 10µM LY display cardiac fusion defects, with the majority of embryos displaying severe cardia bifida phenotypes (Fig. 5E, F).Together these results suggest that PDGF signaling activates PI3K activity to promote myocardial movement towards the midline.

## Discussion

Our studies reveal an intrinsic PI3K-dependent mechanism by which the myocardium moves towards the midline during the formation of the primitive heart tube. Together with our previous studies revealing a role for the PDGF pathway in facilitating communication between the endoderm and myocardium (28), our current work suggests a model in which Pdgfra in the myocardium senses signals (PDGF ligands) from the endoderm and via the PI3K pathway directs myocardial movement towards the midline through the production of medially oriented membrane protrusions. While genetic and imaging studies in zebrafish and mice (5, 13, 18, 19, 25-27, 46, 47) along with embryological studies in chicks and rats (22-24, 48) have identified the importance of extrinsic influences – such as the endoderm and extracellular matrix, on myocardial movement to the midline, our studies using tissue-specific techniques identifies an active role for myocardial cells, providing insight into the balance of intrinsic and extrinsic influences that regulate the collective movement of the myocardial tissue during heart tube formation.

Specifically, we found a requirement for PI3K signaling in cardiac fusion which is complemented by previous studies in mice examining *Pten*, an antagonist of PI3K (29). *Pten* mutant mice also display cardia bifida (33), indicating that appropriate PIP3 levels and localization are required for proper cardiac fusion. Our spatial and temporal experiments further build on these studies by revealing a requirement for PI3K specifically in the myocardium and throughout the duration of cardiac fusion (Fig. 2). We also observed a mild disorganization of the sub-cellular localization of intercellular junctions in the myocardium of PI3K-inhibited embryos (Suppl. Fig. 3). This finding is consistent with previous studies linking epithelial polarity to PI3K signaling (49). However, myocardial cells defective in apical-basal polarity still form a cardiac ring (50, 51), suggesting that an apical-basal defect is unlikely to be the primary reason for myocardial movement defects. Instead, our studies showing that PI3K-inhibited myocardial cells move slower and most prominently are misdirected during the early stages of cardiac fusion indicate a role for PI3K signaling in the steering of myocardial movements medially towards the midline. Our finding that steering in PI3K-inhibited embryos is perturbed in the early stages of cardiac fusion is furthermore consistent with the different phases of myocardial movement identified by Holtzman et al. (9) and suggests that PI3K signaling could be part of a distinct molecular mechanism that drives these early medial phases of myocardial movement. We also found that similar to loss-of-function *pdgfra* mutants, inhibition of PI3K signaling causes defects in directional movement. However, inhibition of PI3K signaling affects myocardial velocity and efficiency (∼ 20% µ/min decrease in velocity, and a 25% decrease in efficiency compared to DMSO-treated embryos) more noticeably than *pdgfra* mutants, in which no significant difference in velocity or efficiency were detected (28). These differences could simply be a result of differences in the extent of PI3K inhibition by 20µM LY compared to extent of loss-of-*pdgfra* function by the *ref* mutation. Alternatively, similar to the role of PI3K signaling in the velocity of gastrulating mesoderm cells as well as in migrating *dictyostelium* and neutrophil cells (30, 31, 36, 52) these differences could also indicate a Pdgfra-independent PI3K function in regulating the velocity of myocardial movement.

Myocardial membrane protrusions were postulated by De Haan et al. ∼50 years ago as a mechanism by which myocardial cells move towards the midline (53). Here using mosaic membrane labeling of myocardial cells to visualize membrane protrusions, we have observed myocardial membrane protrusions that are oriented in the medial direction in a PI3K-dependent manner, confirming his hypothesis. These studies are complemented by previous studies in zebrafish which have observed myocardial protrusions prior to and after cardiac fusion (26, 54) as well as recent studies in the mice (7) indicating that these cellular processes are likely conserved. Indeed, similar observations of PI3K signaling orienting and stimulating protrusion formation in migrating *Dictyostelium* and neutrophil cells as well as in the collective movement of endothelial tip cells, the prechordal plate and the dorsal epithelium (30, 31, 36, 45, 55, 56) support a conserved role for PI3K signaling in regulating protrusion formation.

However, the question of how active membrane protrusions facilitate the collective medial movement of the myocardium to the midline remains to be addressed. Our studies indicate that directionality and to a lesser extent velocity and efficiency are compromised, when membrane protrusions are improperly oriented in PI3K-inhibited embryos (Fig. 3). These observations could suggest that the observed membrane protrusions are force generating, similar to protrusions from leader cells in the lateral line or in endothelial and tracheal tip cells (57-59). Alternatively, these protrusions could act more like filopodia sensing extrinsic signals and the extracellular environment (60). Future studies examining myocardial protrusions and their role in the biomechanical dynamics of the myocardium will help to elucidate the role of membrane protrusions in the collective movement of the myocardium during cardiac fusion.

Overall, our studies delineate a role for the PDGF-PI3K pathway in the mechanisms by which myocardial precursors sense and respond to extracellular signals to move into a position to form the heart. These mechanisms are likely relevant to other organ progenitors including neural crest cells, endothelial precursors, endodermal progenitors, and neuromasts, all of which must move from their location of specification to a different location for organ formation. Although varying in their morphogenesis, many of these movements are collective in nature. Indeed, a similar Pdgfra-PI3K signaling cassette is important in the collective directional migration of several organ progenitors including the migration of mesoderm and neural crest cells (36, 61-67). Receptor tyrosine kinase (RTK)-PI3K pathways are also important across several cardiac developmental processes, including epicardial development, cardiac neural crest addition, cardiomyocyte growth, cardiac fibroblast movement and cardiomyocyte contraction (44, 68-72). Similarly, PDGF-PI3K and more generally RTK-PI3K signaling cassettes are activated in several diseases including glioblastomas, gastrointestinal stromal tumors and cardiac fibrosis (73-76). Thus, the role of this RTK-PI3K cassette in sensing and responding to extracellular signals is likely to be broadly relevant to the etiology of a wide array of developmental processes as well as congenital diseases.

## Supporting information

Timelapse movies

## Acknowledgements

We thank members of the Bloomekatz lab and S. Liljegren, B. Jones, K. Willett, M. Jekabsons, Y. Qiu for helpful discussions; R. Cao, G. Roman, C. Thornton and P. Bolton for imaging and animal support as well as C. Chang, D. Dong, K. Kwan for providing reagents. Funding from the American Heart Association (18CDA34080195) and National institute of Child Health and Human Development (R15HD108782) to JB, and Institutional Development Award (IDeA) from the NIGMS of the NIH (P20GM103460) to the UM GlyCORE imaging facility and JB.

## Contributions

R. Shrestha, T. McCann and J. Bloomekatz conceived of the project and experimental design. R. Shrestha, T. McCann, H. Saravanan, J. Lieberth, P. Koirala and J. Bloomekatz performed experimental studies. R. Shrestha, J. Bloomekatz and T. McCann prepared the manuscript.

## Material Availability

Materials not available commercially are available upon request to Dr. Joshua Bloomekatz.

## Materials and methods

### Zebrafish husbandry, microinjections and plasmid construction

All zebrafish work followed protocols approved by the University of Mississippi IACUC (protocol #21-007). Wildtype embryos were obtained from a mixed zebrafish (*Danio rerio*) AB/TL background. The following transgenic lines of zebrafish were used: *Tg(myl7: eGFP)*^*twu34*^ (RRID:ZFIN_ZDB-GENO-050809-10)(77), *Tg(sox17:eGFP)*^*ha01*^ (ZFIN_ZDB-GENO-080714-2)(78), *Tg(hsp70l:pdgfaa-2A-mCherry;cryaa:CFP)*^*sd44*^ abbreviated *hs:pdgfaa* (ZDB-GENO-170510-4), and *ref (pdgfra*^*sk16*^) (ZDB-GENO-170510-2) (28). All embryos were incubated at 28.5 ºC unless otherwise noted. Transgenic *Tg(myl7:dnPI3K; Cryaa:CFP)* F0 founders were established using standard Tol2-mediated transgenesis (79). F0 founders pairs were screened by intercrosses looking for a high percentage of F1 embryos with CFP+ eyes and cardiac edema. Stable transgenic lines could not be propagated due to loss of viability. Based on the germline mosaicism of the F0 parents, only a proportion of the F1 embryos are expected to have the transgene. Embryos from 4 different F0 pairs were analyzed for cardiac fusion phenotypes. Due to germ-line mosaicism F1 embryos were genotyped after *in situ* hybridization for the presence of the transgene using standard PCR genotyping. Primer sequences are provided in Suppl. Table 1.

Truncated p85 (dnPI3K) capped mRNA was synthesized from the pBSRN3-Δp85 construct (40) and injected at the 1-cell stage. To mosaically label cells in the myocardium for protrusion imaging, *myl7:lck-eGFP* (30 ng/µl) DNA was injected along with Tol2 transposase (40 ng/µl) into *Tg(myl7:eGFP)* heterozgyous embryos at the 1 cell stage and embryos were subsequently allowed to develop at 28.5 ºC.

Plasmids were constructed by using gibson assembly (NEB, E2621) to transfer lck-eGFP (80) or a truncated version of p85 (40) into the middle-entry vector of the tol2 gateway system (81), which were verified by sequencing. Primer sequences are provided in Suppl. Table 1. Then gateway recombination between p5E-myl7 promoter, the constructed middle-entry clones, p3E-polyA and either pDESTTol2pA2 (81) or pDESTTol2pA4-Cryaa:CFP (28) was used to produced plasmids containing *myl7:lck-eGFP* or *myl7:dnPI3K; Cryaa:CFP*, respectively.

### Inhibitor treatments

The following inhibitors were used: LY294002 (LY, Millipore-Sigma 154447-36-6), Dactolisib (Dac, Millipore-Sigma 915019-65-7), and Pictilisib (Pic, Millipore-Sigma 957054-30-7). For each treatment, inhibitors were freshly diluted serially from stocks such that the same percentage (0.1%) of DMSO (Goldbio 67-68-5, Millipore-Sigma 67-68-5) was added to 1X E3 in glass vials (Fisherbrand 03-339-22B). 0.1% DMSO was used as a control. 15 dechorionated embryos per vial were incubated in the dark at 28.5 ºC. In the course of these studies, we noticed that incubation with pharmacological PI3K-inhibitors caused a delay in trunk elongation and somite formation along with defects in cardiac fusion (Suppl. Fig. 2). To ensure our analysis was not obfuscated by a developmental delay, we used somite number to stage match embryos. PI3K-inhibited embryos thus develop approximately 2-3 hours longer than DMSO-treated embryos, prior to analysis.

### Immunoblot, Immunofluorescence, *in situ* hybridization

Embryos at 22s were prepared for immunoblots by deyolking (82). Primary and secondary antibodies include phospho-AKT (1:2000, Cell Signaling 4060, RRID: AB_2315049) and pan-AKT (1:2000, Cell Signaling 4691, RRID: AB_915783), Anti-rabbit HRP-conjugated (1:5000, Cell Signaling 7074, RRID: AB_2099233). pAKT and AKT immunoblots were visualized (Azure 600 Imaging system, Azure Biosystems) and quantified using ImageJ (83) by calculating the ratio of pAkt to Akt. Ratios were normalized to DMSO. To identify *pdgfra/ref* heterozygous and homozygous embryos, embryo trunks were clipped and genotyped as described (28). The body of the embryo including the heart was snapped frozen and stored at -80 ºC. After genotyping, they were pooled via their genotype and analyzed via immunoblot. To activate Pdgfra, embryos expressing the *Tg(hsp701: pdgfaa-2A-mCherry)* transgene were heat-shocked at bud stage as described (28) and collected at 22s.

Immunofluorescence performed on transverse sections used standard cryoprotection, embedding and sectioning (46). Primary, secondary antibodies and dyes include: anti-GFP (1:1000, Abcam ab13970, RRID: AB_300798), anti-ZO-1 (1:200, Thermo Fisher Scientific 33-9100, RRID: AB_87181), donkey anti-chicken-488 (1:300, Thermo Fisher Scientific A32931TR, RRID: AB_2866499), donkey anti-mouse-647 (1:300, Thermo Fisher Scientific A32728, RRID:AB_2633277). TUNEL was performed using the Cell Death detection kit, TMR red (Millipore Sigma 12156792910). Addition of DNaseI was used to confirm we could detect apoptotic cells.

*In situ* hybridization was performed using standard protocols (Alexander et al., 1998), with the following probes: *myl7* (ZDB-GENE-991019–3), *axial* (ZDB-GENE-980526–404) and *hand2* (ZDB-GENE-000511–1). Images were captured with Zeiss Axio Zoom V16 microscope (Zeiss) and processed with ImageJ.

### Fluorescence Imaging

To analyze cardiac fusion (Fig. 1A’-E’) *Tg(myl7:eGFP)* embryos were fixed, manually deyolked and imaged with a Leica SP8X microscope. To analyze the anterior endoderm (Suppl. Fig. 3N-P) *Tg(sox17: eGFP)* embryos were fixed and imaged with an Axio Zoom V16 microscope (Zeiss).

For live imaging, *Tg(myl7:eGFP)* embryos were exposed to DMSO or 20µM LY at bud stage and mounted at 12 somite stage as described (84). Mounted embryos were covered with 0.1% DMSO/20µM LY in Tricaine-E3 solution and imaged using a Leica SP8 X microscope with a HC PL APO 20X/0.75 CS2 objective in a chamber heated to 28.5 ºC. GFP and brightfield stacks were collected approximately every 4 min for 3 hours. After imaging, embryos were removed from the mold and incubated for 24 hrs in E3 media at 28.5 ºC. Only embryos that appeared healthy 24 hours post imaging were used for analysis. The tip of the notochord was used as a reference point to correct embryo drift in the Correct 3D direct ImageJ plugin (85). Embryos were handled similarly for imaging protrusions, except 15 confocal slices of 1µm thickness were collected every 1.5 min with a HC PL APO 40X/1.10 CS2 objective.

### Image analysis

Embryonic length (Suppl. Fig. 2) was measured from the anterior tip of the head to the posterior tip of the tail of each embryo using the free-hand tool of ImageJ. The endoderm width was measured 300 microns anterior from the posterior point of intersection of the two sides of the endoderm. The distance between the *hand2* expressing domains was measured at three equidistant positions (∼200 microns apart) along the anterior posterior axis. *Tg(myl7:eGFP)*+ cardiomyocytes were counted from blinded and non-blinded 3D confocal images of 20s embryos from 4 biological replicates using the cell counter addon in ImageJ. No difference between the blinded (1) and non-blinded (3) replicates was detected.

For live imaging of cell movements – the mTrackJ addon in ImageJ (86) was used. 20-25 cells per embryo whose position could be determined at each timepoint were chosen from the two most medial columns of myocardial cells on each side of the embryos. From these tracks, cell movement properties including overall displacement, velocity (displacement/time), efficiency (displacement/distance) and direction (atan(Δ*y*/Δ*x*)×57.295) were calculated. Rose plots in Fig. 3 display the direction of movement of the overall trajectory of individual cells. In these plots individual cells are grouped into 6 bins based on their net direction of movement; the length of each radial bar represents the percentage of cells in each bin.

For live imaging of myocardial membrane protrusions – stacks were processed in Leica LASX and/or Imaris Viewer (Bitplane) to position the medial edge to the right of the image. Videos of the myocardium were inspected frame by frame in ImageJ for a protrusion. Only cells that were not neighbored by other labeled cells on their medial and lateral edges were analyzed. The direction of protrusion was measured using the “straight line” function a line to draw a line from the bottom of the protrusion to the tip. All protrusions of each cell over the entire recording were measured. Graphs, cartoons and figures were created with Prism (Graphpad), Excel (Microsoft), and Indesign (Adobe).

### Statistics and replication

All statistical analysis was performed in R or Prism (Graphpad). Sample sizes were determined based on prior experience with relevant phenotypes and standards within the zebrafish community. Deviation from the mean is represented as standard error mean or box-whisker plots. In box-whisker plots, the lower and upper ends of the box denote the 25^th^ and 75^th^ percentile, respectively, with a horizontal line denoting the median value and the whiskers indicating the data range. All results were obtained from at least three separate biological replicates, blinded and non-blinded. All replicates are biological. Samples were analyzed before biological sex is determined (87). Raw data and full p-values included in the source file.

## Legends

**Supplemental Fig. 1:**
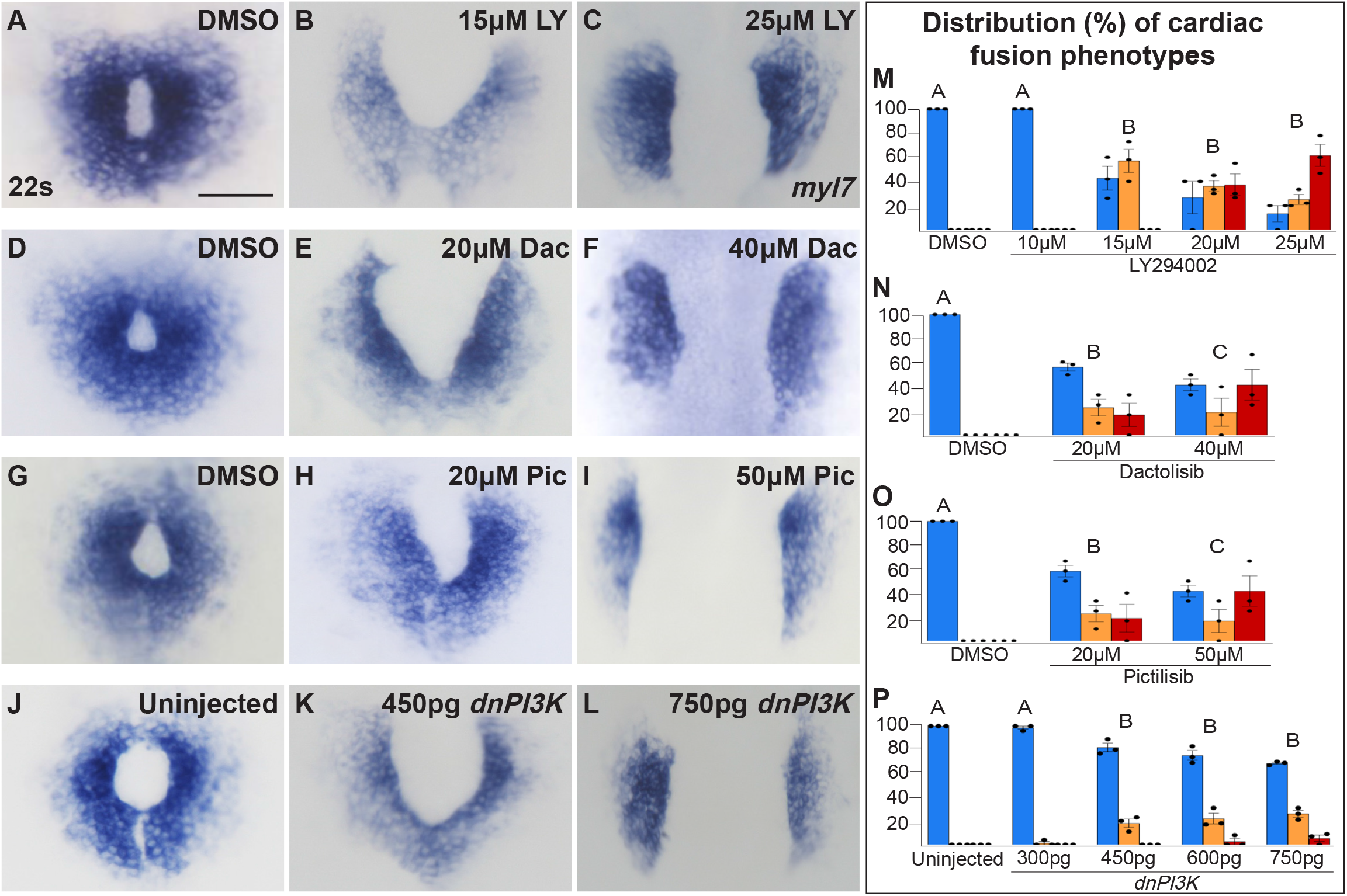
The penetrance and severity of cardiac fusion defects in PI3K-inhibited embryos is dose-dependent. **A-L** Dorsal views, anterior to the top, of the myocardium labeled with *myl7* at 22s. Incubation of embryos with LY (A-C), Dac (D-F), Pic (G-I) from bud stage to 22s or injection of embryos with *dnPI3K* mRNA (J-L) at the one-cell stage results in dose-dependent cardiac fusion defects at 22s. **M-P** Graphs depict the distribution of cardiac fusion defects in embryos treated with increasing concentrations of LY (M), Dac (N), Pic (O) or injected with increasing amounts of *dnPI3K* mRNA (P). Graphs reveal that both the percentage of embryos displaying cardiac fusion defects and the severity of those defects are dose-dependent. Total number of embryos analyzed (n) from > 3 treatments or injections at the indicated concentrations in (M-P): LY-40, 40, 30, 31, 31; Dac: 38, 34, 39; Pic: 37, 39, 38, *dnPI3K* mRNA: 73, 52, 61, 57, 52, respectively. Dots indicate the percent of embryos displaying a specific phenotype per incubation. Blue - Cardiac ring/normal; Orange - fusion only at posterior end/mild, Red - cardia bifida/severe. Bar graphs, mean ± SEM. One-Way ANOVA comparing percent of cardiac fusion defects- letter change indicates p < 0.05. Scale = 60 µm. Full p-values included in the source file.

**Supplemental Fig. 2:**
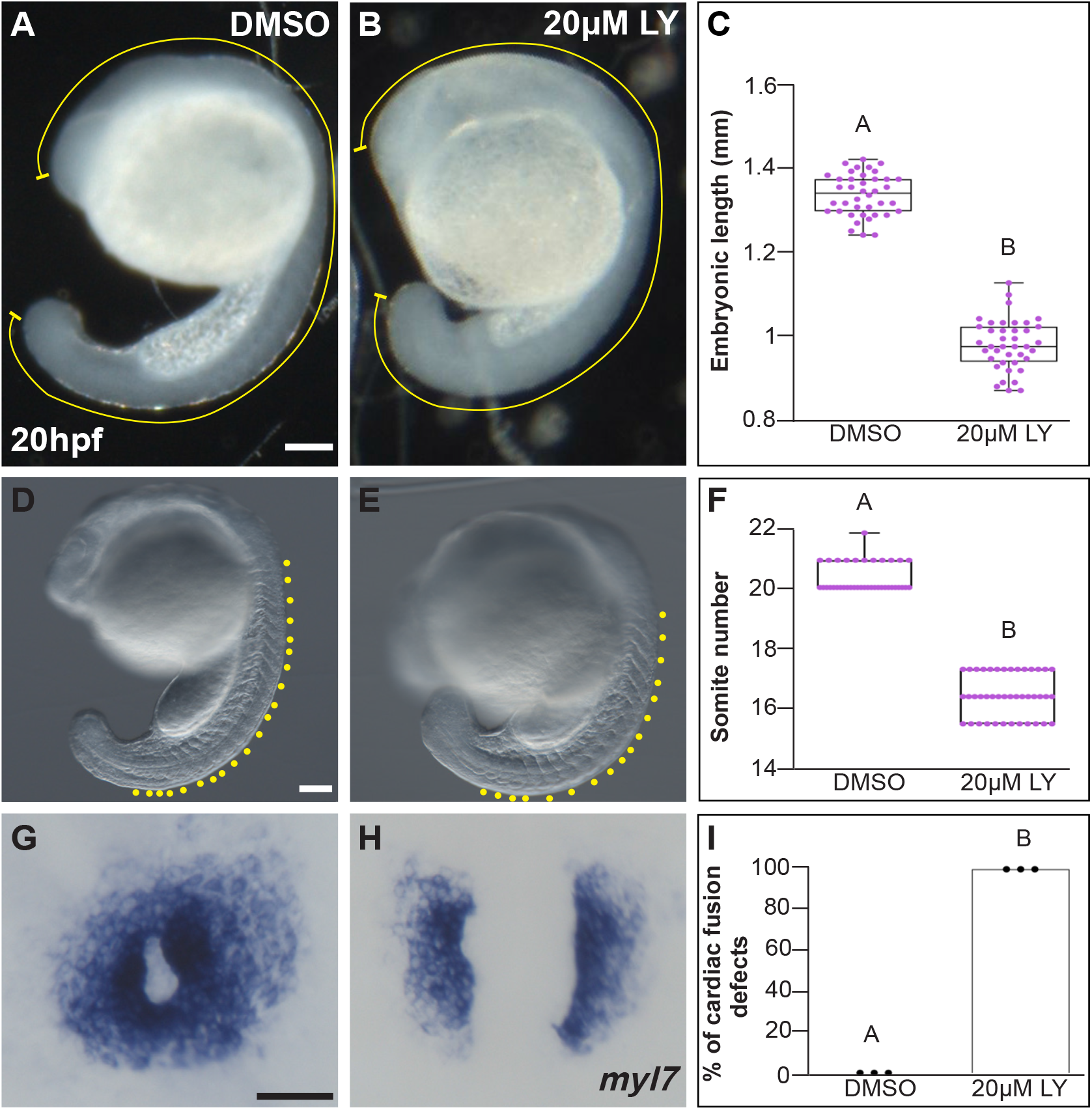
LY-incubation results in trunk extension and somite formation delays. **A-B, D-E** Lateral brightfield views of 20 hours post fertilization (hpf) embryos treated with DMSO (A, D) or 20µM LY (B, E) at bud stage. **C, F** Box-whisker plot depicting the average embryonic length (yellow curved line in A, B) or somite number (yellow dots in D, E) at 20 hpf. Total number of embryos (n) from > 3 separate incubations = 40 (DMSO), 40 (20µM LY) for (C), and 39 (DMSO), 42 (20µM LY) for (F). Dots = measurements from individual embryos. Two sample t-test; p-value = 4.527×10^−4^ and 7.624×10^−5^, respectively. **G-H** Dorsal views, anterior to the top, of the myocardium labeled with *myl7* at 20 hpf. Embryos treated with DMSO at bud stage show cardiac rings (G) whereas those treated with 20µM LY show cardia bifida at 20 hpf (H). **I** Graph depicts the average percentage of cardiac fusion defects in embryos treated with DMSO or 20µM LY. The total number of embryos examined from three separate incubations (n) = 45 (DMSO), 45 (20µM LY). Two sample t-test; p-value = 4.56×10^−5^. Dots indicate the percent of embryos with cardiac fusion defects per incubation. Letter changes (C, F, I) indicate p-values < 0.05. Raw data and full p-values included in the source file.

**Supplemental Fig 3:**
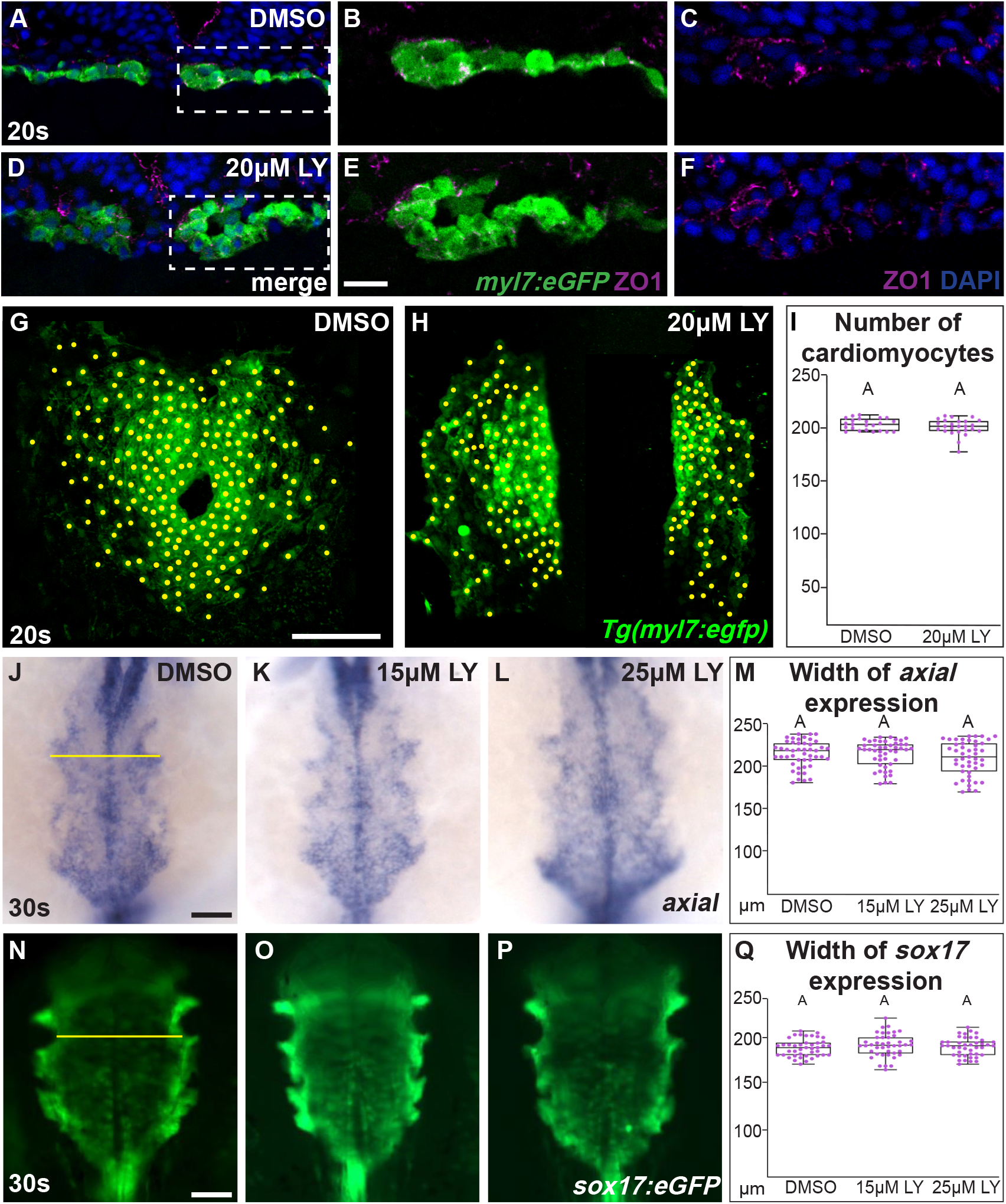
The morphologies of the myocardium and anterior endoderm are not compromised in PI3K-inhibited embryos. **A-F** Representative transverse cryosections, dorsal to the top, compare the morphology of the myocardium, visualized with *Tg(myl7:eGFP)* (green), ZO1 (purple) and DAPI (blue) between DMSO- (A-C) and 20µM LY- (D-F) treated (bud-20s) embryos. Box (A, D) indicate region magnified in B, C, E, F. **G-I** Representative 3D confocal images of the myocardium at 20s, which were used to count myocardial cells in DMSO- (G) or 20µM LY - (H) treated embryos. Yellow dots indicate individual myocardial cells counted using ImageJ. Graph depicts box-whisker plots for the average number of myocardial cells. 21 (DMSO) and 25 (LY) embryos from 4 separate bud-20s incubations were analyzed (I). **J-Q** Dorsal views, anterior to the top, of the anterior endoderm labeled with *axial* (J-L) or the *Tg(sox17:eGFP)* transgene (N-P) at 30s. Embryos incubated with either DMSO (J, N) or 15µM LY (K, O) or 25µM LY (L, P) from the bud stage to 30s show no observable difference in the appearance or width of the anterior endoderm. Box-whisker plots of the average width of the anterior endoderm labeled with either *axial* (M) or *Tg(sox17:eGFP)* (Q). 47 (*axial)* and 42 (*Tg(sox17:egfp)*) embryos per inhibitor concentration from three separate incubations were analyzed. Yellow lines: width of the endodermal sheet. Purple dots (I, M, Q) indicate individual embryos. Letter differences indicate a p-value < 0.05 as tested by 1-way ANOVA. Scale bars, 10 (A-F), 42 (G-H), 60 (J-L), and 50 (N-P) µm. Raw data and full p-values included in the source file.

**Supplemental Fig. 4:**
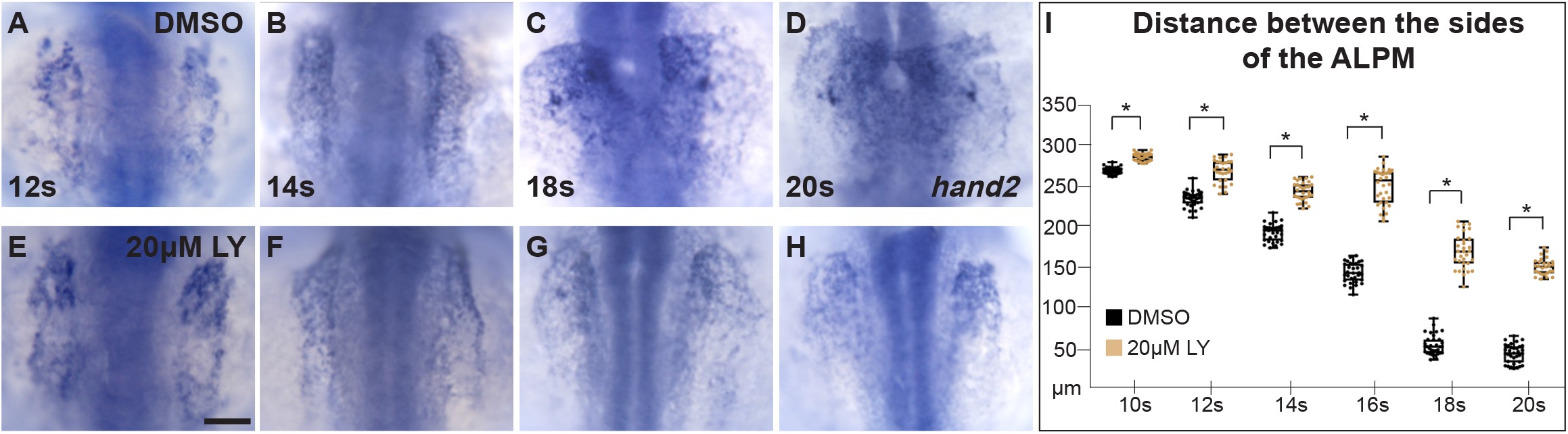
Myocardial movement towards the midline is disrupted in PI3K-inhibited embryos throughout cardiac fusion. **A-H** Dorsal views, anterior to the top, of embryos displaying the expression of *hand2* in the anterior lateral plate mesoderm (ALPM) at (A, E) 12s, (B, F) 14s, (C, G) 18s and (D, H) 20s, treated with either DMSO (A-D) or 20µM LY (E-H) at bud stage. **I** Box-whisker plots depict the average distance between the sides of the ALPM. Although, *hand2* is properly expressed in LY-exposed embryos, ALPM convergence is affected as early as the 10s stage, with a dramatic difference in convergence starting at 12s. The total number of embryos analyzed (n), from 3 separate incubations at the noted stages (I) are: n = 34, 33, 31, 32, 34, 34 (DMSO); 32, 29, 30, 34, 31, 28 (20µM LY), respectively. Dots indicate the distance between ALPM sides per embryo. Student’s t-test: asterisk indicates p values < 0.05. Scale bar, 100µm. Raw data and full p-values included in the source file.

**Supplemental Fig. 5.**
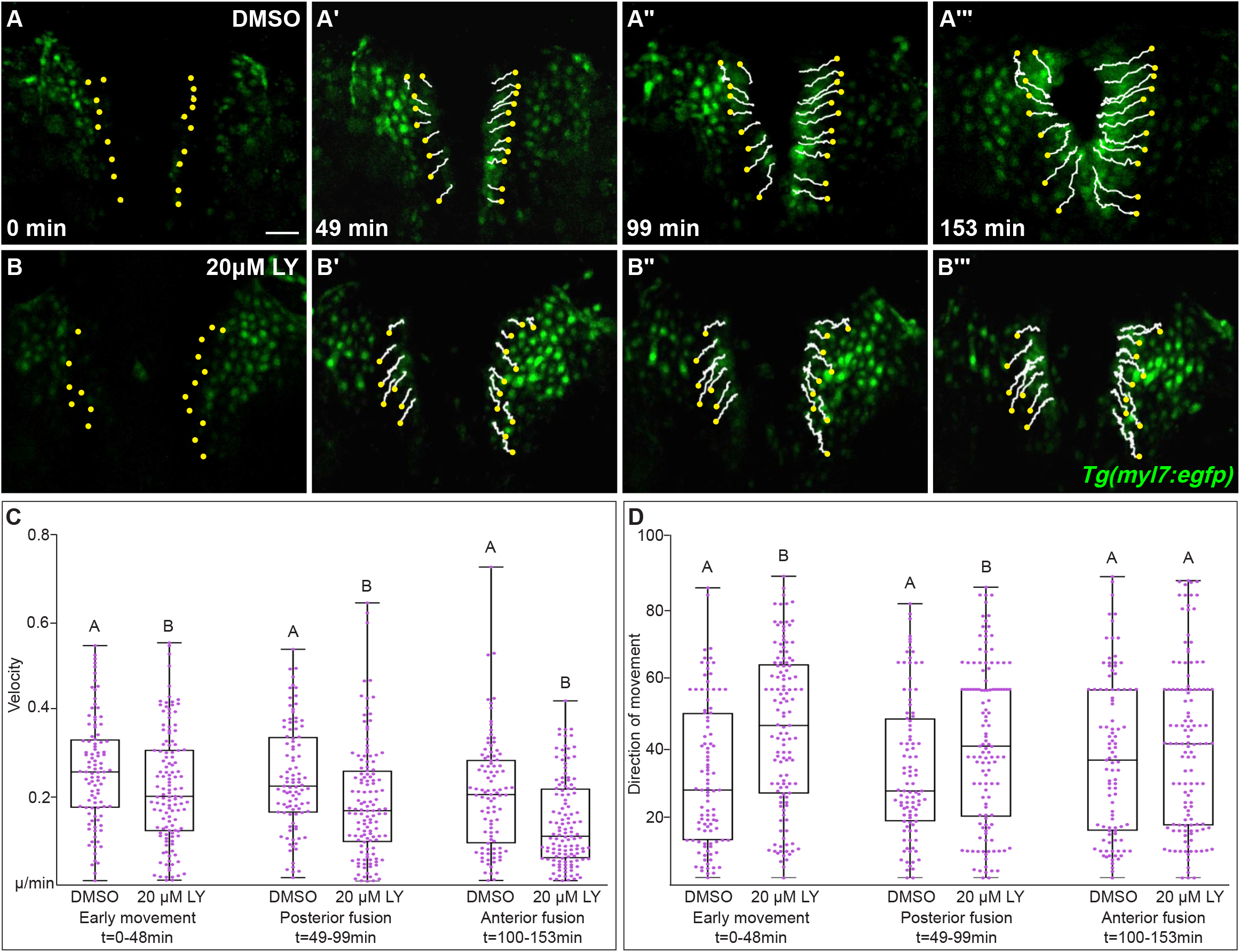
PI3K signaling directs myocardial movement during the early stages of cardiac fusion and regulates velocity throughout cardiac fusion. **A-B** Time-lapse confocal reconstructions from Figure 3 overlaid with cell movement tracks, starting at t = 0 (yellow dots). Scale bar, 60µm. **C, D** Box-whisker plots display the average velocity (C) and direction of movement (D) sub-divided at three distinct stages of cardiac fusion: early movement (0 - 48min), posterior fusion (49 - 99min) and anterior fusion (100 - 153min). The average velocity of myocardial cells in LY-treated embryos is consistently slower than the velocity of DMSO-treated embryos which is consistent throughout cardiac fusion (C). However, in LY-treated embryos myocardial cells display a more angular average direction of movement compared to DMSO-treated embryos during the early stages of cardiac fusion (Early movement, Posterior fusion), after which wild-type myocardial cell movement becomes more angular matching myocardial cell movements in LY-treated embryos. Two sample t-test, letter change indicates p < 0.05. Raw data and full p-values included in the source file.

**Video 1. Myocardial cells in DMSO-treated embryos collectively move towards the midline and form a ring during cardiac fusion. A-C** Representative time-lapse movie of myocardial cells visualized with *Tg(myl7:egfp)* during cardiac fusion in a DMSO-treated embryo (A), tracks show movement of selected cells from the timelapse video (B) and overlay of eGFP and tracks (C). Time-lapse images are of three-dimensional reconstruction of confocal slices taken at 4:32 min intervals for 2.5 hours, beginning at 14s.

**Video 2. Myocardial cells in PI3K-inhibited embryos fail to move properly towards the midline. A-C** Representative time-lapse movie of myocardial cells visualized with *Tg(myl7:egfp)* (A), tracks of selected cells (B), and overlay of tracks and eGFP (C) from an embryo treated with 20µM LY from bud-20s. Time-lapse was acquired as described in Video 1.

**Video 3. PI3K signaling promotes the medial directional movement of myocardial cells towards the midline. A, B** Side-by-side comparison of myocardial movement in DMSO- (A, video 1) and LY- (B, video 2) treated embryos reveals that inhibition of PI3K signaling by LY prevents myocardial cells from being adequately directed towards the midline. Selected analyzed tracks (white lines) overlaying 3D reconstructions of the *Tg(myl7:egfp)* transgene (green) in DMSO (A) and 20µM LY (B) treated embryos.

**Video 4. Dynamic medially oriented myocardial membrane protrusions are lacking in PI3K-inhibited embryos**. Representative time-lapse movies of myocardial membrane protrusions during cardiac fusion, visualized by injecting *myl7:lck-eGFP* plasmids at the 1-cell stage, in DMSO- (left panel) or 20µM LY – (right panel) treated embryos. **Left panel** highlights membrane protrusions (red arrowheads) in a set of posterior myocardial cells in a DMSO-treated embryo. Myocardial membrane protrusions in DMSO-treated embryos are mostly directed in the medial orientation (towards the right in both panels). **Right panel** highlights myocardial membrane protrusions (red arrowheads) in PI3K-inhibited embryos during cardiac fusion. Medial membrane protrusions (towards the right) are lacking in PI3K-inhibited embryos. DMSO or LY treatment from bud-20s. Time-lapse movies are 3D reconstruction of confocal images of membrane protrusions taken at ∼90s intervals for 2 hours. Scale bar, 10µm.

**Supplemental Table S1.**
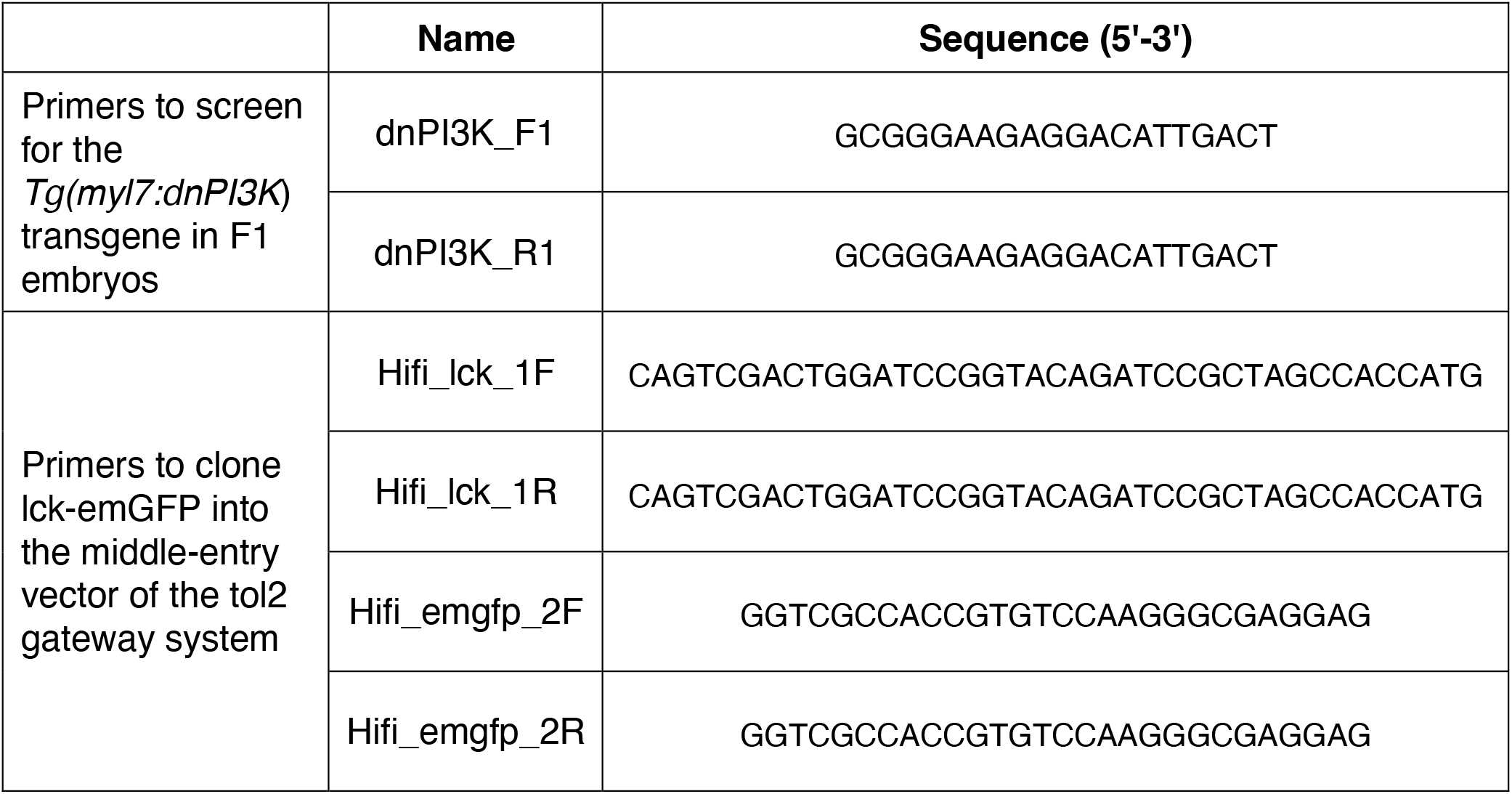
Primers for genotyping and cloning.

## Source data

Excel file organized by figure containing data from which graphs and charts were derived including complete p-values, primer sequences and uncropped immunoblots.

## References

1. Schumacher L. Collective Cell Migration in Development. Adv Exp Med Biol. 2019;1146:105–16.

2. Stainier DY, Lee RK, Fishman MC. Cardiovascular development in the zebrafish. Myocardial fate map and heart tube formation. Development. 1993;119(1):31–40.

3. Wilens S. The migration of heart mesoderm and associated areas in Amblystoma punctatum. Journal of Experimental Zoology. 1955;129(3):579–605.

4. Rawles ME. A study in the localization of organ-forming areas in the chick blastoderm of the head-process stage. Journal of Experimental Zoology. 1936;72(2):271–315.

5. Trinh LA, Stainier DY. Fibronectin regulates epithelial organization during myocardial migration in zebrafish. Dev Cell. 2004;6(3):371–82.

6. Jackson TR, Kim HY, Balakrishnan UL, Stuckenholz C, Davidson LA. Spatiotemporally Controlled Mechanical Cues Drive Progenitor Mesenchymal-to-Epithelial Transition Enabling Proper Heart Formation and Function. Curr Biol. 2017;27(9):1326–35.

7. Dominguez MH, Krup AL, Muncie JM, Bruneau BG. A spatiotemporal gradient of mesoderm assembly governs cell fate and morphogenesis of the early mammalian heart. bioRxiv. y2022:2022.08.01.502159.

8. Linask KK. N-cadherin localization in early heart development and polar expression of Na+,K(+)-ATPase, and integrin during pericardial coelom formation and epithelialization of the differentiating myocardium. Dev Biol. 1992;151(1):213–24.

9. Holtzman NG, Schoenebeck JJ, Tsai HJ, Yelon D. Endocardium is necessary for cardiomyocyte movement during heart tube assembly. Development. 2007;134(12):2379–86.

10. Davidson B, Shi W, Levine M. Uncoupling heart cell specification and migration in the simple chordate Ciona intestinalis. Development. 2005;132(21):4811–8.

11. Evans SM, Yelon D, Conlon FL, Kirby ML. Myocardial lineage development. Circ Res. 2010;107(12):1428–44.

12. Alexander J, Rothenberg M, Henry GL, Stainier DY. casanova plays an early and essential role in endoderm formation in zebrafish. Dev Biol. 1999;215(2):343–57.

13. Kupperman E, An S, Osborne N, Waldron S, Stainier DY. A sphingosine-1-phosphate receptor regulates cell migration during vertebrate heart development. Nature. 2000;406(6792):192–5.

14. Osborne N, Brand-Arzamendi K, Ober EA, Jin SW, Verkade H, Holtzman NG, Yelon D, Stainier DY. The spinster homolog, two of hearts, is required for sphingosine 1-phosphate signaling in zebrafish. Curr Biol. 2008;18(23):1882–8.

15. Kawahara A, Nishi T, Hisano Y, Fukui H, Yamaguchi A, Mochizuki N. The sphingolipid transporter spns2 functions in migration of zebrafish myocardial precursors. Science. 2009;323(5913):524–7.

16. Kikuchi Y, Agathon A, Alexander J, Thisse C, Waldron S, Yelon D, Thisse B, Stainier DY. casanova encodes a novel Sox-related protein necessary and sufficient for early endoderm formation in zebrafish. Genes Dev. 2001;15(12):1493–505.

17. Mendelson K, Lan Y, Hla T, Evans T. Maternal or zygotic sphingosine kinase is required to regulate zebrafish cardiogenesis. Developmental dynamics : an official publication of the American Association of Anatomists. 2015;244(8):948–54.

18. Molkentin JD, Lin Q, Duncan SA, Olson EN. Requirement of the transcription factor GATA4 for heart tube formation and ventral morphogenesis. Genes Dev. 1997;11(8):1061–72.

19. Ye D, Lin F. S1pr2/Galpha13 signaling controls myocardial migration by regulating endoderm convergence. Development. 2013;140(4):789–99.

20. Li S, Zhou D, Lu MM, Morrisey EE. Advanced cardiac morphogenesis does not require heart tube fusion. Science. 2004;305(5690):1619–22.

21. Yelon D, Horne SA, Stainier DY. Restricted expression of cardiac myosin genes reveals regulated aspects of heart tube assembly in zebrafish. Dev Biol. 1999;214(1):23–37.

22. Goss CM. Double hearts produced experimentally in rat embryos. Journal of Experimental Zoology. 1935;72(1):33–49.

23. Rosenquist GC. Cardia bifida in chick embryos: Anterior and posterior defects produced by transplanting tritiated thymidine-labeled grafts medial to the heart-forming regions. Teratology. 1970;3(2):135–42.

24. Varner VD, Taber LA. Not just inductive: a crucial mechanical role for the endoderm during heart tube assembly. Development. 2012;139(9):1680–90.

25. Cui C, Cheuvront TJ, Lansford RD, Moreno-Rodriguez RA, Schultheiss TM, Rongish BJ. Dynamic positional fate map of the primary heart-forming region. Dev Biol. 2009;332(2):212–22.

26. Ye D, Xie H, Hu B, Lin F. Endoderm convergence controls subduction of the myocardial precursors during heart-tube formation. Development. 2015;142(17):2928–40.

27. Aleksandrova A, Czirok A, Kosa E, Galkin O, Cheuvront TJ, Rongish BJ. The endoderm and myocardium join forces to drive early heart tube assembly. Developmental biology. 2015;404(1):40–54.

28. Bloomekatz J, Singh R, Prall OW, Dunn AC, Vaughan M, Loo CS, Harvey RP, Yelon D. Platelet-derived growth factor (PDGF) signaling directs cardiomyocyte movement toward the midline during heart tube assembly. Elife. 2017;6.

29. Fruman DA, Chiu H, Hopkins BD, Bagrodia S, Cantley LC, Abraham RT. The PI3K Pathway in Human Disease. Cell. 2017;170(4):605–35.

30. Iijima M, Devreotes P. Tumor suppressor PTEN mediates sensing of chemoattractant gradients. Cell. 2002;109(5):599–610.

31. Yoo SK, Deng Q, Cavnar PJ, Wu YI, Hahn KM, Huttenlocher A. Differential regulation of protrusion and polarity by PI3K during neutrophil motility in live zebrafish. Dev Cell. 2010;18(2):226–36.

32. Ghiglione C, Jouandin P, Cerezo D, Noselli S. The Drosophila insulin pathway controls Profilin expression and dynamic actin-rich protrusions during collective cell migration. Development. 2018;145(14).

33. Bloomekatz J, Grego-Bessa J, Migeotte I, Anderson KV. Pten regulates collective cell migration during specification of the anterior-posterior axis of the mouse embryo. Dev Biol. 2012;364(2):192–201.

34. Shrestha R, Lieberth J, Tillman S, Natalizio J, Bloomekatz J. Using Zebrafish to Analyze the Genetic and Environmental Etiologies of Congenital Heart Defects. Adv Exp Med Biol. 2020;1236:189–223.

35. Vlahos CJ, Matter WF, Hui KY, Brown RF. A specific inhibitor of phosphatidylinositol 3-kinase, 2-(4-morpholinyl)-8-phenyl-4H-1-benzopyran-4-one (LY294002). J Biol Chem. 1994;269(7):5241–8.

36. Montero JA, Kilian B, Chan J, Bayliss PE, Heisenberg CP. Phosphoinositide 3-kinase is required for process outgrowth and cell polarization of gastrulating mesendodermal cells. Curr Biol. 2003;13(15):1279–89.

37. Gharbi SI, Zvelebil MJ, Shuttleworth SJ, Hancox T, Saghir N, Timms JF, Waterfield MD. Exploring the specificity of the PI3K family inhibitor LY294002. Biochem J. 2007;404(1):15–21.

38. Raynaud FI, Eccles SA, Patel S, Alix S, Box G, Chuckowree I, Folkes A, Gowan S, De Haven Brandon A, Di Stefano F, Hayes A, Henley AT, Lensun L, Pergl-Wilson G, Robson A, Saghir N, Zhyvoloup A, McDonald E, Sheldrake P, Shuttleworth S, Valenti M, Wan NC, Clarke PA, Workman P. Biological properties of potent inhibitors of class I phosphatidylinositide 3-kinases: from PI-103 through PI-540, PI-620 to the oral agent GDC-0941. Mol Cancer Ther. 2009;8(7):1725–38.

39. Folkes AJ, Ahmadi K, Alderton WK, Alix S, Baker SJ, Box G, Chuckowree IS, Clarke PA, Depledge P, Eccles SA, Friedman LS, Hayes A, Hancox TC, Kugendradas A, Lensun L, Moore P, Olivero AG, Pang J, Patel S, Pergl-Wilson GH, Raynaud FI, Robson A, Saghir N, Salphati L, Sohal S, Ultsch MH, Valenti M, Wallweber HJ, Wan NC, Wiesmann C, Workman P, Zhyvoloup A, Zvelebil MJ, Shuttleworth SJ. The identification of 2-(1H-indazol-4-yl)-6-(4-methanesulfonyl-piperazin-1-ylmethyl)-4-morpholin-4-yl-t hieno[3,2-d]pyrimidine (GDC-0941) as a potent, selective, orally bioavailable inhibitor of class I PI3 kinase for the treatment of cancer. J Med Chem. 2008;51(18):5522–32.

40. Carballada R, Yasuo H, Lemaire P. Phosphatidylinositol-3 kinase acts in parallel to the ERK MAP kinase in the FGF pathway during Xenopus mesoderm induction. Development. 2001;128(1):35–44.

41. Alessi DR, Andjelkovic M, Caudwell B, Cron P, Morrice N, Cohen P, Hemmings BA. Mechanism of activation of protein kinase B by insulin and IGF-1. EMBO J. 1996;15(23):6541–51.

42. Osborne N, Brand-Arzamendi K, Ober EA, Jin S-W, Verkade H, Holtzman NG, Yelon D, Stainier DYR. The Spinster Homolog, Two of Hearts, Is Required for Sphingosine 1-Phosphate Signaling in Zebrafish. Current Biology. 2008;18(23):1882–8.

43. Fukui H, Terai K, Nakajima H, Chiba A, Fukuhara S, Mochizuki N. S1P-Yap1 signaling regulates endoderm formation required for cardiac precursor cell migration in zebrafish. Developmental cell. 2014;31(1):128–36.

44. Crackower MA, Oudit GY, Kozieradzki I, Sarao R, Sun H, Sasaki T, Hirsch E, Suzuki A, Shioi T, Irie-Sasaki J, Sah R, Cheng HY, Rybin VO, Lembo G, Fratta L, Oliveira-dos-Santos AJ, Benovic JL, Kahn CR, Izumo S, Steinberg SF, Wymann MP, Backx PH, Penninger JM. Regulation of myocardial contractility and cell size by distinct PI3K-PTEN signaling pathways. Cell. 2002;110(6):737–49.

45. Garlena RA, Lennox AL, Baker LR, Parsons TE, Weinberg SM, Stronach BE. The receptor tyrosine kinase Pvr promotes tissue closure by coordinating corpse removal and epidermal zippering. Development. 2015;142(19):3403–15.

46. Garavito-Aguilar ZV, Riley HE, Yelon D. Hand2 ensures an appropriate environment for cardiac fusion by limiting Fibronectin function. Development. 2010;137(19):3215–20.

47. Arrington CB, Yost HJ. Extra-embryonic syndecan 2 regulates organ primordia migration and fibrillogenesis throughout the zebrafish embryo. Development. 2009;136(18):3143–52.

48. Moreno-Rodriguez RA, Krug EL, Reyes L, Villavicencio L, Mjaatvedt CH, Markwald RR. Bidirectional fusion of the heart-forming fields in the developing chick embryo. Dev Dyn. 2006;235(1):191–202.

49. Krahn MP. Phospholipids of the Plasma Membrane - Regulators or Consequence of Cell Polarity? Front Cell Dev Biol. 2020;8:277.

50. Horne-Badovinac S, Lin D, Waldron S, Schwarz M, Mbamalu G, Pawson T, Jan Y, Stainier DY, Abdelilah-Seyfried S. Positional cloning of heart and soul reveals multiple roles for PKC lambda in zebrafish organogenesis. Curr Biol. 2001;11(19):1492–502.

51. Rohr S, Bit-Avragim N, Abdelilah-Seyfried S. Heart and soul/PRKCi and nagie oko/Mpp5 regulate myocardial coherence and remodeling during cardiac morphogenesis. Development. 2006;133(1):107–15.

52. Hoeller O, Kay RR. Chemotaxis in the absence of PIP3 gradients. Curr Biol. 2007;17(9):813–7.

53. DeHaan RL. Development of form in the embryonic heart. An experimental approach. Circulation. 1967;35(5):821–33.

54. Rohr S, Otten C, Abdelilah-Seyfried S. Asymmetric involution of the myocardial field drives heart tube formation in zebrafish. Circ Res. 2008;102(2):e12–9.

55. Dumortier JG, Martin S, Meyer D, Rosa FM, David NB. Collective mesendoderm migration relies on an intrinsic directionality signal transmitted through cell contacts. Proc Natl Acad Sci U S A. 2012;109(42):16945–50.

56. Graupera M, Guillermet-Guibert J, Foukas LC, Phng LK, Cain RJ, Salpekar A, Pearce W, Meek S, Millan J, Cutillas PR, Smith AJ, Ridley AJ, Ruhrberg C, Gerhardt H, Vanhaesebroeck B. Angiogenesis selectively requires the p110alpha isoform of PI3K to control endothelial cell migration. Nature. 2008;453(7195):662–6.

57. Caussinus E, Colombelli J, Affolter M. Tip-cell migration controls stalk-cell intercalation during Drosophila tracheal tube elongation. Curr Biol. 2008;18(22):1727–34.

58. Dalle Nogare DE, Natesh N, Vishwasrao HD, Shroff H, Chitnis AB. Zebrafish Posterior Lateral Line primordium migration requires interactions between a superficial sheath of motile cells and the skin. Elife. 2020;9.

59. Qin L, Yang D, Yi W, Cao H, Xiao G. Roles of leader and follower cells in collective cell migration. Mol Biol Cell. 2021;32(14):1267–72.

60. Heckman CA, Plummer HK, 3rd. Filopodia as sensors. Cell Signal. 2013;25(11):2298–311.

61. Yang X, Chrisman H, Weijer CJ. PDGF signalling controls the migration of mesoderm cells during chick gastrulation by regulating N-cadherin expression. Development. 2008;135(21):3521–30.

62. McCarthy N, Wetherill L, Lovely CB, Swartz ME, Foroud TM, Eberhart JK. Pdgfra protects against ethanol-induced craniofacial defects in a zebrafish model of FASD. Development. 2013;140(15):3254–65.

63. Bahm I, Barriga EH, Frolov A, Theveneau E, Frankel P, Mayor R. PDGF controls contact inhibition of locomotion by regulating N-cadherin during neural crest migration. Development. 2017;144(13):2456–68.

64. Symes K, Mercola M. Embryonic mesoderm cells spread in response to platelet-derived growth factor and signaling by phosphatidylinositol 3-kinase. Proc Natl Acad Sci U S A. 1996;93(18):9641–4.

65. Klinghoffer RA, Hamilton TG, Hoch R, Soriano P. An allelic series at the PDGFalphaR locus indicates unequal contributions of distinct signaling pathways during development. Dev Cell. 2002;2(1):103–13.

66. He F, Soriano P. A critical role for PDGFRalpha signaling in medial nasal process development. PLoS Genet. 2013;9(9):e1003851.

67. Nagel M, Winklbauer R. PDGF-A suppresses contact inhibition during directional collective cell migration. Development. 2018;145(13).

68. Kim J, Wu Q, Zhang Y, Wiens KM, Huang Y, Rubin N, Shimada H, Handin RI, Chao MY, Tuan TL, Starnes VA, Lien CL. PDGF signaling is required for epicardial function and blood vessel formation in regenerating zebrafish hearts. Proc Natl Acad Sci U S A. 2010;107(40):17206–10.

69. Sato A, Scholl AM, Kuhn EN, Stadt HA, Decker JR, Pegram K, Hutson MR, Kirby ML. FGF8 signaling is chemotactic for cardiac neural crest cells. Dev Biol. 2011;354(1):18–30.

70. Ivey MJ, Kuwabara JT, Riggsbee KL, Tallquist MD. Platelet-derived growth factor receptor-α is essential for cardiac fibroblast survival. American Journal of Physiology-Heart and Circulatory Physiology. 2019;317(2):H330–H44.

71. McMullen JR, Shioi T, Zhang L, Tarnavski O, Sherwood MC, Kang PM, Izumo S. Phosphoinositide 3-kinase(p110alpha) plays a critical role for the induction of physiological, but not pathological, cardiac hypertrophy. Proc Natl Acad Sci U S A. 2003;100(21):12355–60.

72. Shioi T, Kang PM, Douglas PS, Hampe J, Yballe CM, Lawitts J, Cantley LC, Izumo S. The conserved phosphoinositide 3-kinase pathway determines heart size in mice. EMBO J. 2000;19(11):2537–48.

73. Wang SI, Puc J, Li J, Bruce JN, Cairns P, Sidransky D, Parsons R. Somatic mutations of PTEN in glioblastoma multiforme. Cancer Res. 1997;57(19):4183–6.

74. Cheng CK, Fan QW, Weiss WA. PI3K signaling in glioma--animal models and therapeutic challenges. Brain Pathol. 2009;19(1):112–20.

75. Fan H, Ma L, Fan B, Wu J, Yang Z, Wang L. Role of PDGFR-beta/PI3K/AKT signaling pathway in PDGF-BB induced myocardial fibrosis in rats. Am J Transl Res. 2014;6(6):714–23.

76. Lennartsson J, Jelacic T, Linnekin D, Shivakrupa R. Normal and oncogenic forms of the receptor tyrosine kinase kit. Stem cells. 2005;23(1):16–43.

77. Huang CJ, Tu CT, Hsiao CD, Hsieh FJ, Tsai HJ. Germ-line transmission of a myocardium-specific GFP transgene reveals critical regulatory elements in the cardiac myosin light chain 2 promoter of zebrafish. Dev Dyn. 2003;228(1):30–40.

78. Mizoguchi T, Verkade H, Heath JK, Kuroiwa A, Kikuchi Y. Sdf1/Cxcr4 signaling controls the dorsal migration of endodermal cells during zebrafish gastrulation. Development. 2008;135(15):2521–9.

79. Fisher S, Grice EA, Vinton RM, Bessling SL, Urasaki A, Kawakami K, McCallion AS. Evaluating the biological relevance of putative enhancers using Tol2 transposon-mediated transgenesis in zebrafish. Nat Protoc. 2006;1(3):1297–305.

80. Chertkova AO, Mastop M, Postma M, van Bommel N, van der Niet S, Batenburg KL, Joosen L, Gadella TWJ, Okada Y, Goedhart J. Robust and Bright Genetically Encoded Fluorescent Markers for Highlighting Structures and Compartments in Mammalian Cells. bioRxiv. 2017:160374.

81. Kwan KM, Fujimoto E, Grabher C, Mangum BD, Hardy ME, Campbell DS, Parant JM, Yost HJ, Kanki JP, Chien CB. The Tol2kit: a multisite gateway-based construction kit for Tol2 transposon transgenesis constructs. Dev Dyn. 2007;236(11):3088–99.

82. Purushothaman K, Das PP, Presslauer C, Lim TK, Johansen SD, Lin Q, Babiak I. Proteomics Analysis of Early Developmental Stages of Zebrafish Embryos. Int J Mol Sci. 2019;20(24).

83. Stael S, Miller LP, Fernandez-Fernandez AD, Van Breusegem F. Detection of Damage-Activated Metacaspase Activity by Western Blot in Plants. Methods Mol Biol. 2022;2447:127–37.

84. McCann T, Shrestha R, Graham A, Bloomekatz J. Using Live Imaging to Examine Early Cardiac Development in Zebrafish. Methods Mol Biol. 2022;2438:133–45.

85. Parslow A, Cardona A, Bryson-Richardson RJ. Sample drift correction following 4D confocal time-lapse imaging. J Vis Exp. 2014(86).

86. Meijering E, Dzyubachyk O, Smal I. Methods for cell and particle tracking. Methods Enzymol. 2012;504:183–200.

87. Wang X, Bartfai R, Sleptsova-Freidrich I, Orban L. The timing and extent of ‘juvenile ovary’phase are highly variable during zebrafish testis differentiation. Journal of Fish Biology. 2007;70:33–44.

